# Who and how much? Quantifying the role of trophic guilds in soil organic carbon mineralization

**DOI:** 10.64898/2026.04.29.721540

**Authors:** MB Dahl, F Protti-Sánchez, V Groß, A Burns, A Söllinger, D Hu, B Bhattarai, K Dumack, D Metze, I Ostonen, I Janssens, BD Sigurdsson, M Bahn, A Richter, T Urich

## Abstract

Understanding the biotic processes that drive soil organic carbon (SOC) mineralization is essential for predicting the climate warming-carbon cycle feedback. Here, we combined Tree-of-life sequencing (TOLseq; cross-domain profiling using ribosomal RNA) with quantitative conversion factors linking rRNA transcript abundance to biomass, to understand how soil food web changes affect SOC mineralization in an *in situ* soil warming experiment. Field observations showed that warming reduced SOC stocks, but after decades of warming SOC mineralization had acclimated. An energetic soil food web model revealed both bottom-up and top-down controls in the trophic cascades, shifting C flows from fungal and plant-associated channels towards the bacterial channel. This caused an increase in SOC mineralization rate in warmed soils of 30% per-unit biomass across the year.

## Main text

Soil organic carbon (SOC) is a key carbon (C) pool on Earth and essential for soil functions and services including climate regulation (*1*). Recent research has identified organic molecules derived from microbial necromass as main components of SOC and emphasized the so-called ‘microbial carbon pump’, which conceptualizes both how C is dynamically recycled within microbial biomass and channelled into long term storage in the soil (*2, 3*). The microbial pump is controlled by predation and other trophic interactions in the soil food web (*4*), and potentially also by virus-induced lytic events (*5, 6*). Consequently, SOC mineralization is not carried out by single species, but rather a result of biotic interactions of an unmatched diversity of organisms (*7*). The comprehensive assessment of the soil food web (SFW) required to understand these C dynamics is hampered by the difficulty in extractability, spatial heterogeneity, enormous diversity and temporal variation of soil (micro)biomes (Fig. 1A), rendering it *de facto* still not possible (*8, 9*). Modern molecular techniques such as *T*ree-*O*f-*L*ife sequencing (TOLseq; metatranscriptomic sequencing of ribosomal RNA for three-domain profiling, (*10, 11*)) provide the methodological approach allowing to capture the entire SFW structure in one analysis. However, utilizing the information-rich data derived from such ‘omics’ techniques to predict ecosystem responses to e.g., climate change, remains a key challenge in the field of ecosystem science (*12*). Recent research has emphasized how the application of energetic food webs (formulated decades ago (*13–16*)) has the potential to close the gap between soil biodiversity and C flow (biology and biogeochemistry). Food web approaches inherently encompass multiple trophic levels by estimating C flow through trophic networks (i.e., in a predation matrix). However, community data are often labour-intensive to collect and tend to focus on higher trophic-level chains, leaving microorganisms poorly resolved at best (*17, 18*).

**Fig. 1.**
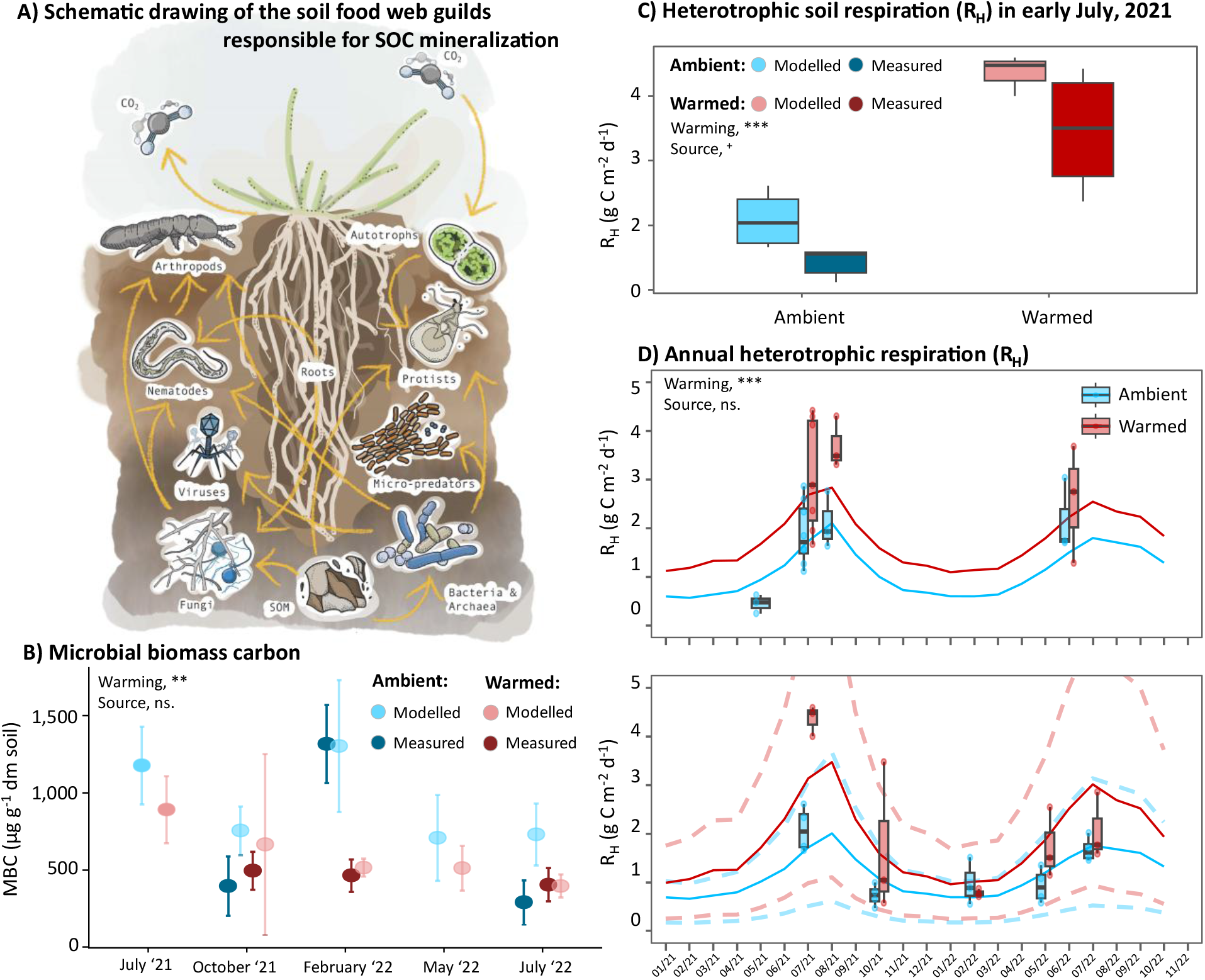
Soil food web modelled and measured heterotrophic soil respiration. (**A**) Schematic overview of the studied soil food web organisms. (**B**) Measured and modelled microbial biomass carbon (MBC) at each season and for ambient and warmed soil temperature conditions. (**C**) Direct comparison of the soil heterotrophic respiration (RH) for early July 2021 (measured on July 7^th^ and 14^th^ 2021) to modelled rates (based on TOLseq profiles from July 14^th^ 2021). (**D**) Soil heterotrophic respiration (RH) extrapolated with Van’t hoff equation for annual dynamics of modelled and measured rates (measured rates are summarized per month). Confidence intervals (broken lines) on the modelled rates represents the 80% CI of the multifactorial sensitivity test of a half-to-double variable range (see Material and Methods and Fig. S8). *** p < 0.001, ** p < 0.05, ^+^ p = 0.06, ns = non-significant linear mixed-effects model.

Here, we demonstrate how quantitative TOLseq profiling of the soil (micro-)biota — combined with easily measurable variables such as microbial biomass C and standard physiological parameters — can serve as a foundation for constructing energetic food web models. This approach enables us to disentangle the structural drivers of food web dynamics underlying observed bulk biogeochemical processes, a methodology we term the TOLmodel. The novel approach was applied here on soil samples originating from unmanaged grassland sites in Iceland (‘Forhot’ sites (*19*)), where geothermal activity over a period of more than 60 years created the soil warming gradients of up to +6 °C used in this study (for details see Material and Methods). Our TOLmodel approach allowed the quantification of soil biota and their contributions to SOC mineralization rates, and uncovered seasonal patterns across trophic groups and under different warming intensities.

### From rRNA to Biomass

It has been demonstrated that energetic food webs can be parameterized using only community data (i.e., abundance of organisms in a given ecosystem), with remaining parameters inferred from the literature (*17*). Quantitative soil metatranscriptomics, which considers the total extracted RNA from a sample (*20*), has emerged as a key technology for linking microbiome abundance and activity to biogeochemical processes. Applying this method, studies have shown that transcript mass (µg transcripts g^-1^ sample) — unlike relative transcript abundance — can reveal dynamic changes in community size and transcriptional activity, which align with gas flux measurements, as demonstrated for the temporal dynamics of methane emission and methanogens in different soils (*21, 22*) and cow rumen (*20*).

However, linking rRNA transcripts to the biomass of organisms has remained a challenge. As molecular assessments, such as DNA or RNA sequencing, are becoming increasingly used, factors converting these molecules’ abundances into biomass are emerging. For example, it was recently suggested that extracted soil DNA related to microbial biomass carbon (MBC) across different soil types by a multiplication factor of 15 (for sand and silt loams (*23*)). However, when relying on nucleic acid (NA) extractions to reflect absolute abundance, one needs to consider the NA capture efficiency from the soil matrix, which varies with soil type (*23–26*). One way to overcome this problem is to include an internal extraction standard during NA extraction (*27, 28*), such as an RNA standard to obtain quantitative estimates of ribosomal RNA transcripts in the soil (*29*).

Ribosomes represent a considerable fraction of cell compounds (*30, 31*). Their mass scales to cell size, although the ratio varies with intrinsic (e.g., growth phase, (*32*)), environmental (e.g., temperature (*33*)) and/or nutrient conditions (*34*). For this reason, we here constrained the total ratio of rRNA to biomass by quantifying both rRNA (*20,29*) and MBC (chloroform fumigation). The ribosome-to-biomass ratio also differs among organisms. In *Escherichia coli*, ribosomal RNA account for 20% of dry mass (dm) when grown under optimal laboratory conditions (*35*, and BNID 104954, *36*), while for yeast (*S. cerevisiae*) the proportion was shown to be 5-10% (BNID 108196, (*37)*). This relationship of approx. 1:3 is supported by our laboratory observations from pure cultures (Fig. S1). While only a few studies exist for metazoa, a study on different species of freshwater arthropods showed that RNA represented between 3-6% of dm (*38*). Based on the above studies, we here assumed that the ribosome-to-biomass ratio scales 1:3:4 for prokaryotes, microbial eukaryotes, and multicellular organisms, respectively.

Applying this approach, in our study system the ribosomal RNA in prokaryotes were found to represent 5.9 ± 4.1 % of the total dm-C. Since bacterial members of the SFW in the present study represented a broad range of sizes and life strategies (Fig. S2), and the majority of the bacterial population (up to 60% (*39)*) would likely be in a starved or only ‘potentially active’ state at any given time point (i.e. with fewer ribosomes), the estimated ribosomes-to-biomass ratio was considered reasonable. The ratio was observed to change according to temperature with a potential Q10 of 0.7 (albeit a poor regression fit, Fig. S3). A similar temperature sensitivity was observed for microbial turnover (Q10 = 0.8, R^2^ = 0.4, Fig. S4). Since ribosomes are key cellular machinery needed for protein biosynthesis and growth, it seems likely that ribosomes would be tightly correlated to and show a similar temperature sensitivity as turnover rates. Furthermore, it is in agreement with an increasing understanding of how ribosomes are tightly regulated according to environmental conditions (*33*).

The average standing MBC, estimated from the measured rRNA, was significantly lower in warmed soils than in ambient soils (600 ± 220 and 930 ± 260 µg MBC g^-1^ dm soil, respectively, p = 0.004, Fig. 1B), in line with previous observations for MBC from chloroform fumigated measurements at the same site (*40*). Especially during winter, a large difference in MBC between ambient and warmed soils was observed (significant for both measurements): 1,320 (± 250) and 460 (± 110) µg biomass-C g^-1^ dm soil, and ribosomal-based estimates of 1,300 (± 420) vs. 510 (± 60) µg MBC g^−2^ dm soil.

### Realistic estimation of SOC mineralization rates by TOLmodel

From TOLseq profiles (1.14 × 10^8^ SSU rRNAs, an average of 2.8 ± 0.5 million sequences per sample, n = 35, Data S1), trophic entities of the SFW were classified. The unprecedented taxonomic resolution of bacteria, archaea, fungi, protists, animals and viruses, enabled via TOLseq, was sub-structured into a total of 34 functional guilds of the SFW (Data S2 and exemplified in Fig. 1A). Laboratory measurements of microbial turnover and previously reported microbial carbon use efficiency (CUE) from the sites (*40*), was used to parameterize the speed and efficiency of C transfer in the food web (Material and Methods, Table S1).

Modelled mineralization rates were compared to *in situ* soil CO2 efflux measurements from root exclusion collars, which represent heterotrophic soil respiration (RH), a close proxy of SOC mineralization. Measurements of RH were conducted between May and September of 2021 and 2022. A direct comparison between modelled and measured rates was possible for days in early July, 2021 (Material and Methods). Modelled and measured values aligned (although linear mixed-effects model revealed a marginally significant difference, p = 0.06) and soil warming significantly increased RH for both measured (from 1.6 ± 0.6 for ambient to 3.5 ± 1.0 g C m^-2^ d^-1^ for warmed, p < 0.001, Fig. 1C) and modelled (2.1 ± 0.5 to 4.3 ± 0.3 g C m^-2^ d^-1^, p < 0.001) and showed a temperature sensitivity corresponding to a Q10 between 1.9 to 2.6 (measured) and 2.2 to 2.4 (modelled, Fig. S5). By using RH rates at mean annual soil temperature annual RH rates were estimated (*41, 42*), which were 360 ± 10 g C m^−2^ yr^−1^ (measured) and 380 ± 10 (modelled) for ambient, and 610 ± 50 g C m^−2^ yr^−1^ (measured) and 530 ± 40 g C m^−2^ yr^−1^ (modelled) for warmed soils (Fig. 1C). These rates are consistent with values reported in the literature, where average RH in ‘temperate grassland’ soils was approximately 300 g C m^−2^ y^−2^ (*43, 44*). A sensitivity analysis found that the model parameters ‘production efficiency’ and ‘ribosome-to-biomass ratio’ had the strongest effect on model outputs (Fig. S6, S7)—particularly for *Prokaryota*, which accounted for the majority of the biomass in the studied system (Fig. 2A–B). A multi-factorial sensitivity analysis (i.e., considering combinations of variability) was used to infer a confidence interval for the modelled SOC mineralization rates (dashed lines in Fig. 1C). The multifactorial analysis found that 87.6 ± 9.6% of the predicted RH rates fell within the value range of measured RH (Fig. S8). Predictions falling outside the bounds of the field observations were primarily due to unrealistic parameter combinations—for example, the convergence of low biomass-to-ribosome ratio (high biomass estimate) and low production efficiency at high turnover rates (e.g., ‘Summer’, Fig. S8). Such scenarios necessitated implausibly high C fluxes through the organisms to maintain steady-state biomasses under the given food web structure, or *vice versa* low biomass-to-ribosome ratio (low biomass estimate) and high efficiency at low turnover rates resulting in minimal mineralization (e.g., ‘Winter’ and ‘Autumn’, Fig. S8).

**Fig. 2.**
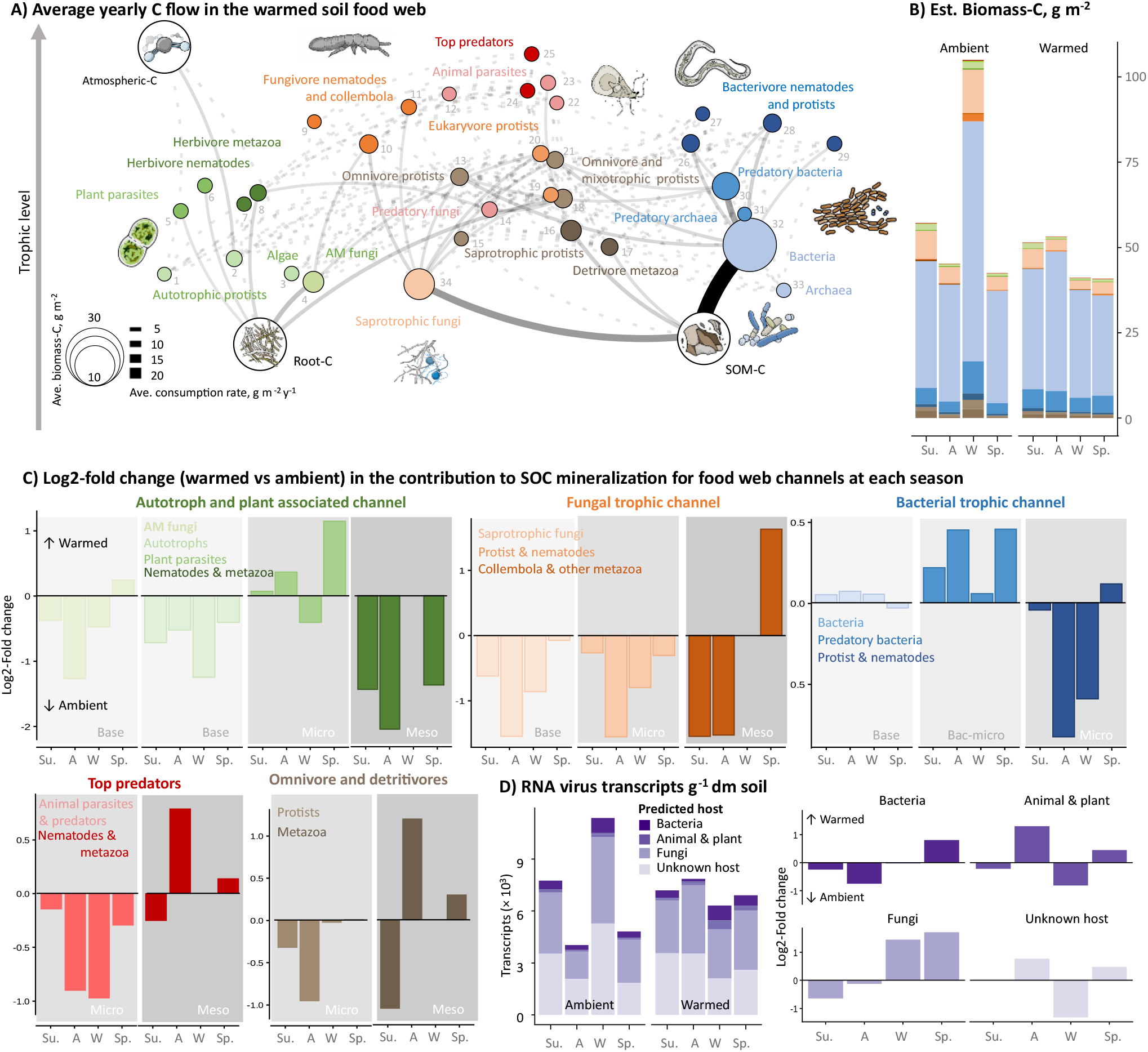
Soil food web composition and contributions to SOC mineralization by trophic modules. (**A**) Schematic overview of C flow in the SFW. Broken lines represent C flows < 0.1 g m^-2^ y^-1^. Members of each node can be by node number in Data S3. (**B**) Estimated biomass of trophic groups, g biomass-C m^-2^ per warming condition and season. (**C**) The log2-fold change in contributions to SOC mineralization for each trophic group between ambient and warmed conditions per season. (**D**) Transcripts of RNA viruses (normalized to 1 million reads per sample) classified by predicted host under ambient and warmed conditions for each season. Su.: Summer (averaged for 2021 and 2022), A: Autumn, W: Winter, Sp.: Spring. Background grey-scale indicate the size of organisms, distinguishing ‘micro’ and ‘meso’ next to the ‘base’-level of each trophic chain, exception for predatory bacteria denoted ‘bac-micro’ acknowledging them as the smallest predator in the food web.

### Quantifying the impact of soil food web topology on SOC mineralization

With the community profiles coupled to masses and mineralization rates (Fig. 1B–D) individual contributions to SOC mineralization could be identified for each trophic guild identified in the food web. Saprotrophic bacteria accounted for the largest proportion of SOC mineralization across all seasons and under both temperature conditions (dominated by e.g., *Acido*-, *Actino*- and *Proteobacteria*, 84 ± 2% of C mineralization, Table S2), which is in line with theoretical expectations (*45*). Hereafter followed, in ambient soils, saprotrophic fungi (e.g., *Helotiales* and *Archaeorhizomycetales*, with an annual average contribution of 6.6 ± 1.0%), whereas in warmed soils predatory bacteria (e.g., *Myxococcota*) contributed 7.4 ± 0.3 % and saprotrophic fungi only 4.0 ± 1.2% (Table S2). These differences at the base-level of the SFW had cascading effects in the trophic structure, where the entire fungal C-channel was reduced under warmed conditions (log2FC < 0, with one exception in spring of mesofauna, e.g., *Collembola*, orange panel in Fig. 2C). A similar pattern was seen for the plant-associated C channel, where especially plant feeding mesofauna (e.g., *Aphididae* and *Delphacidae*) were reduced with up to 4-fold under warmed conditions (log2FC = -2, green panel Fig. 2C) and only plant parasites were increased under warmed conditions, likely further adding to the negative effect on the plant-associated channel here. Contrary, in the bacterial channel a reduction at higher trophic-levels (nematodes and protists; blue and brown panel in Fig. 2C, respectively) co-occurred with an increase at the base- and first-trophic level, suggesting a release from top-down control by predation on SOC mineralization here.

The reduction in mesofauna in warmed soils is in line with the previously observed trophic-downgrading (i.e. loss of larger soil animals) of the SFW at the warmed sites (*46, 47*). It was previously hypothesized that the lower standing pool of SOC and MBC, together with a higher bulk density, indirectly selected against larger soil animals (*46*). The present study reveals that the lower food resources in the warmed soils reduced the abundance of metazoa in the fungal and plant-associated channels (bottom-up control), whereas general microbivorous metazoa (e.g., *Rotifera*) and omnivores (e.g., *Annelida*; *Lumbricidae* and *Enchytraeidae*) as well as the metazoan top predators (e.g., *Mesostigmata*, Data S3) were not affected (Fig. 2C).

The observed shift towards C cycling at the lower trophic-levels likely has consequences for microbial necromass formation, which is increasingly identified as key for long-term SOC storage (*48,49*), and one that depends on the ‘death pathway’ (*50*). An indirect indication of increased necromass formation under warming was shown in microbial N acquisition, which had shifted towards recycling organic-N from microbial residues (*51,52*). Finally, dynamics in the viral community may further fuel the faster turnover under warming by lysis events, although these are still poorly understood (*6, 53*). Warming did not significantly affect bulk viral RNA, which followed the standing biomass, with significantly more viral transcripts present during winter under ambient conditions, in line with the higher standing biomass at this time (Fig. 1A, 2B and S9). Among the predicted host groups, RNA viruses associated with fungi were the most abundant (Fig. 2D).

When normalizing to rRNA-based MBC, the estimated SOC mineralization rates in the warmed soils were consistently doubled across season (log2-fold change of approx. 1, Fig. S10A). This is in line with the previously reported increase in both biomass-specific microbial growth, respiration and organic-C uptake but an unchanged CUE from the warmed sites (*40*). Long-term warming has increased prokaryotic β-diversity as a result of species composition turnover with a threshold at 4 °C for warm-adapted species (*54*). When screening the prokaryotic taxa in the present study for a genetic adaptation towards faster growth (using the ribosomal rRNA operon copy numbers as a proxy) no indication of a phylogenetic shift was seen (Fig. S11). Despite reducing the standing microbial biomass, warming was shown to increase bacterial richness, i.e., the number of active taxa (*55*), a relationship which was also observed in our TOLseq profiles (Fig. S12). Although warming itself accounted for the majority of the effect, the rates after adjusting to a base temperature of 10 °C (from a Q10 of 2.3, Fig. S5), were still consistently 30% higher under warmed conditions (log2-fold change of approx. 0.35, Fig. S10B). A possible explanation is the shift in food web structure. In ambient soils, the larger fungal channel transfers more C to higher trophic levels (soil animals, mainly *Collembola*), whose longer life span and larger body sizes buffer the SOC mineralization rate by immobilizing C (*4, 48*) and/or by transforming SOC to less degradable forms (*56*). In contrast, under warming the food web channels more C through small single-celled organism and micro-predators (mainly predatory bacteria, *Myxococcota*), whose faster growth and death rate could accelerate SOC mineralization rates. A shift in root-derived C inputs (lower fine-root biomass, *57*) and changes in root exudates (*58*) likely contributed to the reduced fungal abundance under long-term warmed conditions.

The effect size of the changes in trophic structures were quantified as changes in the base-level contribution to SOC mineralization (here saprotrophic bacteria and fungi) in response to the abundance of higher trophic groups. These relationships strongly decreased as predators got bigger (Table 1). For example, a theoretical 10-fold increase in predatory bacteria, would result in an absolute increase of approximately 20 µg C mg^-1^ biomass m^-2^ d^-1^ at 10°C for saprotrophic bacteria, which represents a 67% increase relative to the current average contribution (30 ± 5 µg C mg^-1^ biomass m^-2^ d^-1^ at 10°C; Table S2). Contrary, a 10-fold increase in fungal metazoan grazer abundance corresponded to an absolute increase of 7 µg C mg^-1^ biomass m^-2^ d^-1^ at 10°C from fungi, representing a 24% increase. That is, despite the minor direct contribution to SOC mineralization by the organisms of higher trophic levels (0.2 ± 0.1 % of all mesofauna under ambient, Table S2), the abundance of soil fauna in the trophic chain indirectly determines mineralization rates, suggesting that soil fauna represents an important top-down control on the SOC mineralization.

**Table 1.** Change in the per biomass contribution to SOC mineralization at base-level by 10-fold increase in the abundance (biomass) of higher trophic levels. Only significant correlations are summarized and the R^2^ is given for the equation: y = a × ln(x) +b; where y is the contribution to SOC mineralization by base-level in µg C mg^-1^ biomass m^-2^ d^-1^ and x is the abundance of the higher trophic level (ln(mg m^-2^)). x10 effect was calculated as a × ln(10). Contribution rates are normalized to 10°C. Graphic summary available in Fig. S13.

| Basal trophic level | Higher trophic level | x10 effect | $R^2$ | P-value |
| --- | --- | --- | --- | --- |
| Saprotrophic bacteria |  |  |  |  |
| Direct relationships | Predatory bacteria, bac-micro | 19.7 | 0.35 | < 0.001 |
|  | Bacterivore, micro | 9.7 | 0.16 | 0.03 |
|  | Omnivore, micro | (8.5) | (0.12) | (0.058) |
|  | Omnivore, meso | (7.5) | (0.23) | (0.070) |
| Indirect relationship | Plant feeders, micro | 8.5 | 0.32 | < 0.001 |
| Saprotrophic fungi |  |  |  |  |
| Direct relationship | Fungal feeders, meso | 7.11 | 0.42 | < 0.001 |
| Indirect relationship | Animal, top predators | (3.73) | (0.11) | (0.067) |

### Model application to other grassland soil ecosystems

To further validate TOLmodel, we applied it to three previously published TOLseq datasets from temperate grassland soils (*21, 22*), where parallel *in situ* bulk soil respiration had been measured. These datasets represented ecosystems with relatively low (< 1.0 g C m^-2^ d^-1^, ‘cattle grazed’) and high (up to 6.8 g C m^-2^ d^-1^, ‘drained peatlands’) SOC mineralization rates. In the present study, RH accounted for approximately 65% of bulk soil respiration; we assumed the same proportion for the external datasets. Furthermore, the Q10 response in turnover rates (0.8) and ribosome-to-biomass scaling factor (0.7) was directly transferred without project specific fitting. Given the lack of an internal nucleic acid extraction standard in these studies, no extraction correction was applied. Despite these constraints, TOLmodel yielded SOC mineralization rates that aligned well with the RH estimates from measured soil respiration across all three tested ecosystems (R^2^ = 0.8, p < 0.001, Fig. 3). With the exception of Grassland I — where an outlier prevented a reliable linear regression — and also considering some outliers for Grassland III, it is notable that also for the individual grasslands the regressions approximated a one-to-one relationship, with only a slight positive bias in the intercept (0.5 and 1.3, for Grassland II and III respectively) and a slope close to unity (0.9 for both).

**Fig. 3.**
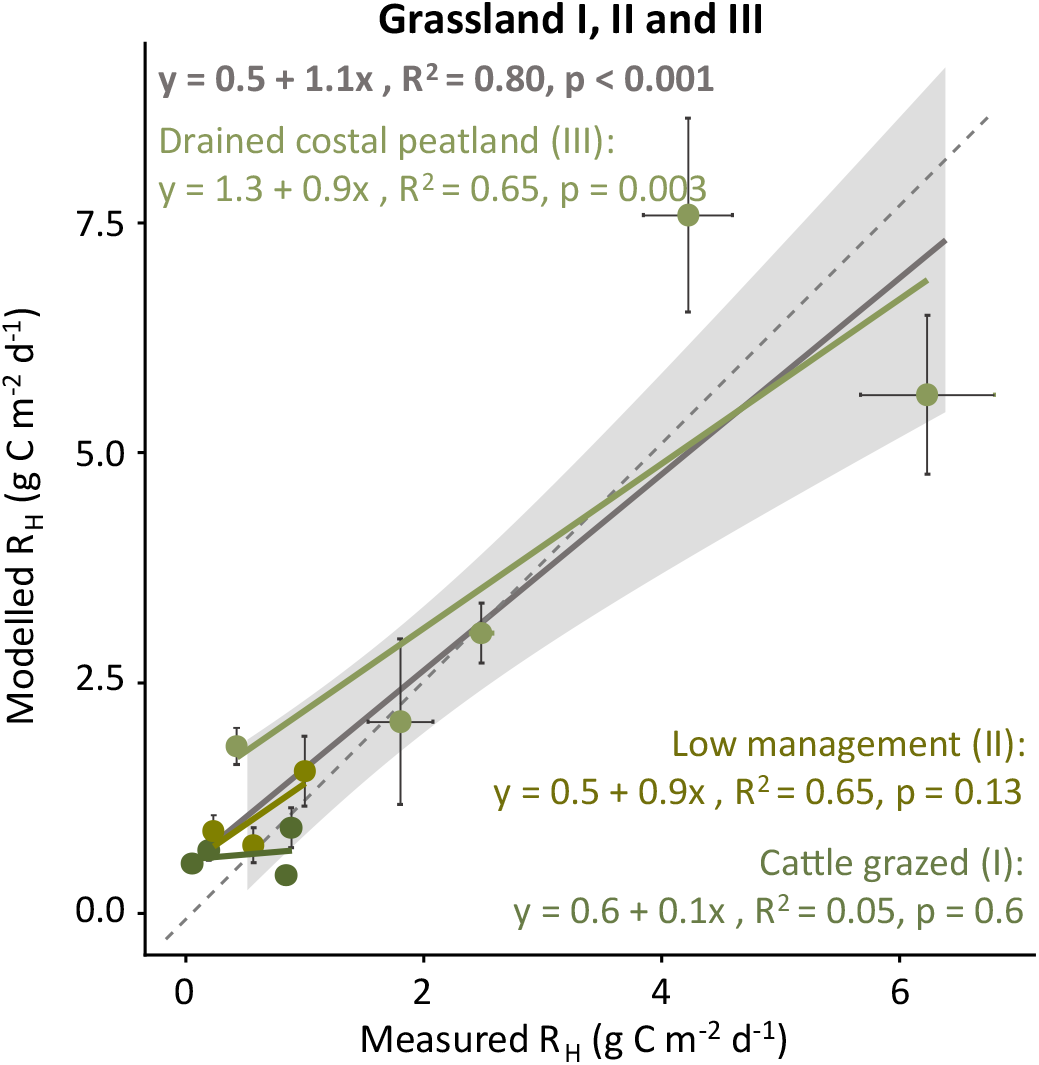
Measured and modelled rates of heterotrophic soil respiration (RH) in three grasslands differing in SOC mineralization. Measured RH refers to estimates based on *in situ* soil respiration measurements. Grey regression represents a combined analysis with 95% confidence intervals (grey shade). Error bars represent S.E. for modelled (vertical) and measured (horizontal) values, replication was available for; three timepoints for Grassland III and one timepoint for Grassland II. For the latter the resulting S.E. was 0.04 and thus not visible. For ‘Grassland III’, a single non-replicated datapoint was considered an outlier and was discarded (model and measured value of 13.8 and 3.2 g C m^-2^ d^-1^, respectively). The dashed line represents the 1:1 line.

Also here, *Bacteria* were identified as the main contributor to SOC mineralization in all grassland soils, albeit with clear seasonal variation (Table S3). In Grassland I and II saprotrophic fungi were the second highest contributors (6 ± 8%), followed by predatory bacteria (5 ± 5%). While in Grassland III omnivore (6 ± 6%) and bacterivore protists (3 ± 3%) were among the main contributors and equally important to predatory bacteria (4 ± 4%).

## Conclusions

Our analyses show that integrating standard measurements of microbial biomass and physiology with quantitative rRNA transcript abundance (TOLseq) bridges molecular ‘omics’ approaches with biochemical process monitoring. This integration enables the identification of key influences of the soil food web topology on SOC mineralization—insights that cannot be inferred from molecular relative abundance data or bulk process measurements alone. We reveal how both bottom-up and top-down trophic cascades shape the food web under long-term warming: the decline of large mesofauna stems primarily from disruptions at the base of the food web (bottom-up), while shifts in intermediate trophic levels — particularly bacterial-feeding protists and nematodes — reduce predatory control within the bacterial C channel, leading to accelerated SOC mineralization.

The TOLmodel framework paves the way for quantifying the effect size of distinct SFW configurations on SOC mineralization. As trait databases continue to grow (*59–61*), improved model parameterization will become increasingly feasible, unlocking the full potential of this integrative approach for understanding ecosystem C dynamics.

## Acknowledgement

We wish to thank Pall Sigurðsson and Coline Le Noir de Carlan for collecting soil samples, Stella Brachmann for preparing metatranscriptomic libraries, Christopher Gall for measuring soil chemicals and microbial parameters as part of their respective MSc thesis’s and Erik Verbruggen for fruitful discussions. We thank Prof. Joanne Emerson and Dr. Luke Hillary for expert assistance with the viral transcript bioinformatics. We are thankful to the editors and reviewers for their valuable suggestions and comments. University available AI tools (OpenWeb UI) were used occasionally to improve wording.

## Funding

This work was funded by the following: The German Research Foundation (DFG) grant BO 5559/1-1 (433256088) and UR198/7-1. ‘FutureArctic’, a European Union’s Horizon 2020 research and innovation program under the Marie Skłodowska-Curie Actions (grant no. 813114). AS acknowledges funding from the Research Council of Norway (FRIPRO project SHRINK, 344999). IO was supported by Estonian Research Council grants PRG916, PRG2614 and by CoE FutureScapes (TK232). AR acknowledges funding from the Austrian Science Fund (FWF grant DOI 10.55776/COE7).

## Author contribution

Conceptualization: MBD and TU. Fieldwork: MBD, AS, DM and BDS. RNA extraction and metatranscriptomic library preparations: VG and students. Laboratory measurements of soil DOC, TC, water content, microbial Cmic, Nmic: AR and students. *In situ* CO2 flux measurements: FPS and MB. Plant root biomass data: BB and IO. Literature collection on SFW predators: VG. Expert knowledge on protist predators: KD. Model testing: MBD and AB. Visualization: DM and MBD. Funding acquisition: BDS, IJ, AR, TU and MBD. Writing original draft: MBD and TU. All authors contributed to the final manuscript.

## Competing interests

the authors declare no conflict of interest.

## Data and materials availability

Data used are described in detail in the SM. Sequencing files are available from NCBI (BioProject: RJNA1099624).

## Supplementary Materials

Material and Methods

Figs. S1 to S15

Tables S1 to S4

Data S1 to S4

## Notes

### Competing Interest Statement

The authors have declared no competing interest.

