## Supplements for "Who and how much? Quantifying the role of trophic guilds in soil organic carbon mineralization"

Dahl *et al.* 2026

**The PDF file includes:**

Material and Methods

Figs. S1 to S15

Tables S1 to S4

**Other Supplementary Materials for this manuscript include the following:**

Data S1 to S4

#### List of content

Material and Methods……......................................................................................... 3

### Material and Methods

#### *Sites and samples*

In this study, all samples originate from natural grassland sites in Iceland, which are part of the 'ForHot' experiment (www.forhot.is and (*62*)). The grassland site consists of replicated soil temperature gradients formed by natural geothermal warming of the bedrock below the soil profile. The site has experienced warming for more than 60 years and is denoted long-term warming (LTW). Samples were collected from four replicated transects with elevated (LTW-$E_T$; +6 °C above ambient) and ambient (LTW-$A_T$) soil temperature; however, samples from transect 3 (LTW-3-$E_T$) were excluded from the analysis because cooling had occurred in the warmed plots in this transect since 2017 (Fig. S14). Soil cores (0-10 cm layer; approx. 40 g) were collected from each plot at each site and immediately frozen on dry ice or in liquid nitrogen. Soil sampling was carried out on 14$^{th}$ July 2021, 26$^{th}$ October 2021, 15$^{th}$ February 2022, 20$^{th}$ May 2022 and 26$^{th}$ of July 2022 (n = 35).

The long-term warming has led to a loss of soil organic matter (SOM) from ca. 35 t C $ha^{-1}$ to 15 t C $ha^{-1}$ and increased the soil bulk density from ca. 0.6 g $cm^{-3}$ to 0.8 g $cm^{-3}$ in LTW-$E_T$ compared to LTW-$A_T$ (*63*). Fine root biomass was quantified in 2019; a total of 30 soil cores (ø = 4.8 cm) were sampled and separated into two soil depths: topsoil (0–10 cm) and subsoil (10–30 cm). Samples were thoroughly washed in the laboratory using a 1-mm sieve over white plastic boxes. All below-ground plant organs were inspected under a dissecting microscope, and separated into living and dead fractions based on texture, consistency, and color (following (*64*)). Only the living fraction was retained for subsequent analyses. Samples were oven-dried at 50 °C for 96 hours to achieve a constant dry weight. Belowground plant root biomass was calculated per ground surface area (g $m^{-2}$) and averaged for in LTW-$E_T$ and LTW-$A_T$ for the food web model (166 and 74 mg $g^{-1}$ soil for LTW-$A_T$ and LTW-$E_T$, respectively, Data S4).

#### In situ *soil carbon efflux measurements*

Heterotrophic soil respiration ($R_H$) was measured in PVC root-exclusion collars (Ø 10 cm), inserted to >20 cm depth, i.e., beyond the site rooting zone (0–10 cm; (*63, 65*)). This approach severs roots and prevents root ingrowth, thereby removing the autotrophic component of soil respiration (*66*). Collars were installed in summer 2018 in LTW-$E_T$ and LTW-$A_T$ plots (n=4). $R_H$ was measured in several campaigns over 2021 and 2022 using a portable infrared gas analyser (SCR-1, EGM-4, PP-Systems, UK) and a closed chamber placed air-tight on the collars for over 2 minutes.

Because the site is affected by geogenic $CO_2$ degassing (6*7*), the biogenic and geogenic $CO_2$ contributions were differentiated using a stable carbon isotope approach. In June 2022, sequential headspace sampling from a chamber (height 16 cm, Ø 10 cm) placed on the collars for 30 min were performed. Gas was sampled at 1, 5, 10, 20 and 30 min (15 mL transferred into 12 mL pre-evacuated Labco Extainer® glass vials) and analysed for $CO_2$ concentration and $\delta^{13}C$ in a gasbench-IRMS (Finnigan MAT). For each series, the isotopic composition of the total soil $CO_2$ was estimated using the Keeling plot approach (*68*). The contribution of geogenic $CO_2$ efflux was quantified using a two-pool mixing model with δ13C end-member values of −2.5‰ (geogenic source, measured directly in gas from nearby vents) and −28‰ (biogenic source, characteristic of C3 plants, (*67*)). Across plots, geogenic effluxes were 0.17 ± 0.06 μmol $CO_2$ $m^{-2}$ $s^{-1}$ (LTW-$A_T$) and 1.91 ± 1.57 μmol $CO_2$ $m^{-2}$ $s^{-1}$ (LTW-$E_T$). These plot-specific geogenic $CO_2$ effluxes were used to correct later campaigns measurements of the total soil $CO_2$ efflux.

#### Microbial biomass carbon (MBC) and turnover

As previously reported, microbial biomass carbon (MBC) was measured from chloroform fumigation extracts. In brief, 15 ml 1 M KCl was added to an aliquot of 2 g of fresh soil per sample and placed on an orbital shaker for 30 min at 200 rpm. A second aliquot of 2 g of fresh soil was fumigated with chloroform ($CHCl_3$) for 48 hours in the dark and then extracted with KCl as described above. Extracts were subsequently filtered through ash-free filter paper (Satorius 392 filters, Göttingen, Germany) and frozen at -20°C until analysed for extractable organic carbon (DOC) on a TOC analyzer (Shimadzu TOC-L, TNM-L). Microbial biomass carbon (MBC) was calculated as the difference between DOC content in the fumigated and non-fumigated extracts. MBC were corrected for incomplete extraction following standard corrections of 0.45 (*69*). To reach microbial biomass estimates, carbon was assumed to represent 50% of the total dry weight biomass (*70*).

Microbial biomass turnover was determined by the $^{18}O$ water vapor equilibration methods (*71*), where the synthesis of new DNA is estimated from the incorporation of $^{18}O$ in genomic DNA. In brief, 0.5 g fresh soil in a 1.0 ml extraction vial was incubated for 48 hours in a 27 ml air-tight glass vials with 500 μL of $^{18}O$-enriched water at the bottom of larger vials (without direct contact to the soil) so that a total enrichment (soil water and external water) was 25 at% $^{18}O$. Subsequently, the $^{18}O$ enrichment of the soil water was determined in a parallel incubation where water samples were taken at the start of the incubation, after 4, 16, 24 and 48 hours (*71*). DNA was extracted from the soils after 48h (MP Biomedicals following manufacturer's protocol,). DNA concentrations were quantified fluorometrically using the PicoGreen Assay (Quant-iTTM PicoGreen dsDNA Reagent, Life Technologies) on a TECAN Infinite 200 PRO plate reader. Isotopic composition of the DNA extracts was analysed on a thermochemical elemental analyzer (TC/EA, Thermo Fisher Scientific) coupled via a Conflo III open split system (Thermo Fisher Scientific) to an IRMS (Delta V Advantage, Thermo Fisher Scientific). Turnover time was calculated as the ratio of total DNA content (ng $g^{-1}$ dw soil) to DNA production rate (ng $g^{-1}$ dw soil $h^{-1}$), normalized by incubation time.

#### *Sequence libraries and bioinformatics*

Mechanical soil homogenization was performed using a Retsch MM 400 mixer mill (Retsch GmbH, Haan, Germany) while keeping the sample (approx. 40 g fresh weight per plot) and containers frozen with liquid nitrogen. The resulting soil powder was stored at -70°C. RNA was extracted from each soil sample (of ca. 0.5 g soil powder) using The RNeasy® PowerSoil® Total RNA Kit (Qiagen, Venlo, Netherlands). Extractions were done following manufacturer's protocol, but with the following modifications: each soil sample was subjected to two cycles of bead-beating (for lysing and homogenization) with FastPrep-24™ 5G (MP Biomedicals, Irvine, CA, USA) to maximize the recovery of RNA per sample, combined directly after phase separation in a joint capture tube. Furthermore, each sample was spiked with 30 ng of purified RNA of a nucleic acid extraction standard ($NAE_{std}$, based on *Saccharolobus solfataricus* (*29*)). The spiking (pipetting and a brief reverting of the tube) was done immediately after the homogenization and prior to the first phase separation during NA extraction. Adding the $NAE_{std}$ directly to the soil-buffer-slurry mixture aims at exposing the $NAE_{std}$ to the same soil chemistry as the released RNA from the soil community (for details see (*29*)). RNA extract was cleaned using column purification with MEGAclear™ Transcription Clean-Up Kit (Thermo Fisher Scientific, Waltham, MA, USA). Finally, the RNA extract (50 µl) was mixed with 1.25 µl RNasin (1 µM) and stored at -70 °C for later use, or subjected to quantitative and qualitative analysis. RNA concentration was determined with Qubit™ RNA BR Assay Kit (Thermo Fisher Scientific, Waltham, MA, USA). RNA quality was analysed by Bioanalyzer 2100 using the Agilent RNA 6000 Nano Kit (Agilent Technologies, Santa Clara, CA, USA).

##### *Metatranscriptomic library preparation*

Libraries were prepared using NEBNext® Ultra™ II RNA Library Prep Kit for Illumina® (New England BioLabs, Ipswich, MA, USA) following manufacturer's protocol. RNA input was 100 – 120 ng. RNA was fragmented to obtain the target fragment length of 370 bp (250 bp insert size + 120 bp for adaptor/primer) for paired-end sequencing. Size selection was performed with HighPrep PCR beads (MagBio Genomics Inc., Gaithersburg, MD, USA). The libraries were paired-end sequenced using a NextSeq 550 System using the NextSeq 500/550 High Output Kit v2.5 (300 Cycles; Illumina, San Diego, CA, USA) at the sequence facility at Greifswald University. Raw sequencing files are available from NCBI (BioProject: RJNA1099624).

##### *Bioinformatic NGS-data processing*

Raw sequences were submitted to the 'PhyloFLASH' processing pipeline following default settings (*72*). In brief, the pipeline identifies SSU rRNA sequences by aligning sequences to a modified SILVA v.138 (NR99; (*73*)) database file using 'BBmap' mapping algorithm (*74*), all matches with a minimum identity of 70% (default) is kept and taxonomic affiliation is assigned by the Last Common Ancestor (LCA) approach. For *Bacteria* and *Archaea*, the LCA taxonomy obtained from 'PhyloFLASH' was directly carried over to the data analysis, whereas all sequences assigned to *Eukaryota* were extracted from the raw reads and submitted to the following assembly pipeline; SSU rRNA euk-reads were pooled for all samples and assembled using SPAdes *(75)* with 97, 107, 117 k-mers and multi-cell mode and annotated against the EUKARYOME SSU database v.2 (*76*) using QIIME2 (*77*) and LCA assigned based on 50 best matches. Length of contrigs (Fig. S15). Reads were mapped back to generate abundance using Bowtie2 *(78)*.

Putative viral reads were identified by first sorting the sequences into rRNA and non-rRNA fractions using SortMeRNA (*79*). The non-rRNA reads were pooled and assembled using Megahit v1.2.9 (*80*). Identification and taxonomy classification of RNA viruses were performed using RdRpCATCH (https://github.com/dimitris-karapliafis/RdRpCATCH) which includes eight different RNA-dependent RNA polymerases (RdRp) Hidden Markov Model (HMM) databases. The potential host of RNA viruses were predicted by RNAVirHost (*81*). A total of 834 potential viral contigs were identified as RNA viruses assigned to 19 orders and six predicted hosts were identified and assigned to their best match. Sample-specific non-rRNA reads were mapped to the RNA viral contigs using Bowtie2 v2.4.1 (*78*), and their abundances were normalized to the total number of annotated non-rRNA reads per sample.

#### *Data analysis and statistics*

All statistical and graphic analyses were conducted in 'R' v.4.3.1 (R Core Team 2024) using mainly the packages *vegan* (*82*), *ggplot2* (*83*), *reshape2* (*84*), *soilfoodweb* (*85*) and *phyloseq* (*86*).

##### *Biomass estimates from RNA sequencing profiles*

From the relative abundance of the $NAE_{std}$ in the SSU rRNA community profiles, the average RNA extraction efficiency was determined to 18.2 ± 13.9 % (Data S4). Total number of ribosomal transcripts per sample were calculated following the quantitative metatranscriptomics approach (qMetra in (*20*, *29*)). In brief, a theoretical ratio of 4:96 for mRNA:rRNA is considered for the total RNA per sample (*87*), which was in agreement with the empirical determined 2.5 ± 1.0% of non-rRNA sequences for the dataset (Data S4). Furthermore, the SSU rRNA is considered $\frac{1}{3}$ of the total rRNA weight (*88*). Hereafter follows to estimate how many rRNA transcripts this represents. The length of 16S and 18S SSU rRNA was considered to be 1,500 and 1,900 nt respectively, and the molecular weight of ssRNA as: No. nucleotides × 320.5 Da + 159 Da accounting for 5'triphosphate per nucleotide; 1 Da = 1 g $mol^{-1}$. In combination with the Avogadro constant of ca. $6.02 \times 10^{23}$ units per mol, we estimated 16S and 18S rRNA transcripts $g^{-1}$ dm soil. For the mass of individual ribosomes we distinguished between prokaryotes (2.5 MDa based on *E. coli*; BNID 106864, *37*), microbial eukaryotes (3.3 MDa for *S. cerevisiae*; BNID 110369, *37*) and multicellular eukaryotes, for the latter we adapted the following adjustment: we consider that eukaryotic ribosomes have increased in molecular mass throughout evolution by the accumulation of rRNA expansion segments and additional ribosomal proteins, with mammalian ribosomes being as large as 4.2 MDa (BNID 106865, *37*). Thus, for the multicellular eukaryotes, which are metazoans (e.g. small soil arthropods such as springtails and mites) but lack vertebrate-level translational complexity (*89*), an intermediate value of 3.7MDa was assumed.

The ribosome-to-biomass relationship was defined as:

$$Biomass = a \times P + 3a \times E + 4a \times M \qquad \text{Eq. 1}$$

Where, **a** is the scaling factor representing the ribosomes-to-biomass ratio for prokaryotes. **P** is the total weight of prokaryotic ribosomes, **E** the same for microbial eukaryotic and **M** for multicellular organism (e.g. soil arthropods). The relationship between the scaling factor for prokaryotes, microbial eukaryotes, and multicellular organisms were based on literature examples (discussed in the main text).

MBC was used as a proxy for microbial biomass, which was assumed to constitute 50% of total biomass (*15*, *70*, *90*). Assuming that mesofauna biomass-C is likely not captured by chloroform fumigation, the scaling factor *a* was determined as:

$$MBC = a \times P + 3\,a \times E \qquad \text{Eq. 2}$$

A variation in the calculated scaling factor ‘a’ depending on temperature was observed (Fig. S3) and the ribosome-to-biomass ratio was thus adjusted according to soil temperature at the time of sampling, with a Van't Hoff equation:

$$\frac{1}{RiboF_Y \times Q10^{\frac{T_X - 10}{10}}} \qquad \text{Eq. 3}$$

Where $RiboF_Y$ is the scaling factor (‘a’ from Eq. 2) for the organism $type_Y$ (prokaryotes, single- or multicellular eukaryotes), $T_x$ is the soil temperature of the $sample_X$ and $Q_{10}$ was 0.7 informed from $RiboF_{Prokaryota}$ (Fig. S3).

*Genetic adaptation of growth rate to warming*

Prokaryotic taxa were screen in the genome-database rrnDB (version 2025-04-07, available here: https://rrndb.umms.med.umich.edu/) and the community-weighted mean of rRNA operon copy numbers per sample was calculated to determine adaptation towards faster growth under LTW-E conditions. We included species-, genus- and family-level annotations. At family-level approximately 42% of the prokaryotic community was annotated, with no significant difference in coverage between plot types, 41.4

± 0.7% and 42.6 ± 0.8% for LTW-E and LTW-A. For neither family nor genus-level annotations was a significant difference in community weighted abundance of rrn between plot types was seen (2.9 ± 0.1 and 2.7 ± 01 cwm-rrn for ambient and warmed plots respectively, ANOVA, $p = 0.62$, Fig. S11).

##### *Energetic soil food web model – TOLmodel parameterization*

Taxonomically annotated SSU rRNA TOLseq profiles were functionally classified according to literature and expert knowledge. For the two major groups of protists *Amoebozoa* and *Rhizaria* we used the recently published trait-database (*91*) to establish feeding mode. Metazoa were classified following Potapov et al. 2022 (*92*) supplemented with a few specific publications (*93–95*). The classification resulted in 34 distinct food web entities (interaction matrix is available in Data S2).

Predation pressure was weighted according to relative abundance (*85*). Besides the interaction (feeding) matrix and the quantified (biomass) abundance table, the model relies on the following parameters: i) assimilation(a) and ii) production efficiency (p), adapted from recent application and original publication on soil food web modelling (*86*, *47*, *15*, *13*), for the latter we adapted the carbon-use-efficiency (CUE) determined for microorganims from the site (0.22, *40*). Similarly, laboratory measurement of microbial biomass turnover (see above and Fig. S4) was used as iii) death rate (d; inverse turnover time given in days), for multicellular eukaryotes 365 days was applied. All parameters are available form Table S1.

Furthermore, at sub-zero temperatures multicellular organisms were considered dormant (i.e., not representing a predation pressure in the food web) and their biomass abundance was set to near-zero and their process variables were set to have no impact on the system ($d = 0.01$, $a = 1$, $p = 1$, as recommended, *85*). Energetic soil food web (SFW) model was run using *comana()* available from the R package *soilfoodwebs* (*85*).

Similarly, we evaluated the per biomass estimated mineralization rates for the food web structure under ambient and warmed conditions across seasons, and for this corrected for temperature as following:

$$\left. \frac{Est.Cmin.}{Est.Biomass} \middle/ Q10^{\frac{T_x - 10}{10}} \right. \qquad \text{Eq. 4}$$

Where $T_x$ is the soil temperature of a given sample and $Q_{10}$ was set to 2.3 (informed form *in situ* measurements and the modelled C flux, Fig. S5).

##### *Sensitivity test of model parameters*

We combine taxa abundance profiles obtained from TOLseq with generalized standard variables of cell physiology and stoichiometry. The original authors (*96*) showed how some of these variables are sensitive to generalization across taxa, i.e., in an analysis of the impact of generalizing across one or more trophic groups. The authors showed how especially the assimilation (A) and production (P) efficiency resulted in the largest error rate for mineralisation estimates when food webs were large (> 20 nodes) and when merging across food web entities happened at higher trophic levels. Similarly, the authors found the death rate and biomass to increase the error of the estimated SOC mineralization when organism at higher trophic levels with a different diet were combined.

Here, we performed a two-step sensitivity analysis of our model parameterization. First, all 16 parameters were tested in a single-factor sensitivity test, considering a variability range of half to double (1,000 permutations). Note that for 'assimilation' and 'production' efficiencies, the upper bound was limited to 1. Test outputs were used to identify influential variables, by random forest approach with the R package *randomForest* (*97*) (Fig. S7). From this evaluation, it was evident that five parameters were the main determiners for the model output, namely i) assimilation efficiency, ii) the production efficiency, and iii) the ribosome-to-biomass ratio (Fig. S7 and main text). Thus, in a subsequent multi-factorial sensitivity analysis (i.e., considering variation in a combinations of multiple parameterizations) was performed in 1,000 permutations of random co-assignment of values in the tested range of values (Fig. S8). Test outputs from the multi-factorial sensitivity test were used to determine a quasi-80% confidence interval for the estimated SOC mineralization (reported in Fig. 1C).

##### *Direct comparison to* in situ *heterotrophic soil respiration measurements*

Measurements of heterotrophic soil respiration were conducted between May and September of 2021 and 2022. Parallel TOLseq sampling and gas flux measurements were conducted on July 14, 2021. However, after correction for the geogenic $CO_2$ efflux, only one $CO_2$ flux replicate from warmed soils was retained for July 14. Therefore, measurements from July 7 (when weather conditions were comparable, Table S4) were included in the directcomparison. Measured and modelled heterotrophic soil respiration was evaluated using a linear mixed-effects model (R package *nlme*, (*98*)), with warming level (LTW-$E_T$ and LTW-$A_T$), Source (modelled or measured), and their interaction as fixed effects, and transect as a random effect. Fixed effects were tested with t-tests using containment approximations for denominator degrees of freedom, as implemented in *nlme*. Pairwise comparisons of estimated marginal means were performed on the fitted model (post-hoc test) with Tukey's adjustment for multiple testing (R package *emmeans*, (*99*)).

##### *Application for previously published datasets*

To evaluate the performance of the model in other soil ecosystem three previously published metatranscriptomic datasets was used. All datasets originated from German grassland soils and had parallel *in situ* measurements of bulk soil respiration ($CO_2$ efflux) available (*21*, *22*). All model settings remained the same and the Q10 response of the ribosome-to-biomass and to turnover rates were adapted

(0.7 and 0.8, respectively), and C flux was calculated as described above. When available, replicated TOLseq profiles (and gas measurements) from the same sample site were used to infer variation for modelled (and measured) C fluxes.

*Supplementary Figures S1-S15 and Table S1-S4*

*Fig. S1 – RNA-to-biomass relationship for a soil bacterial and fungal yeast species.*

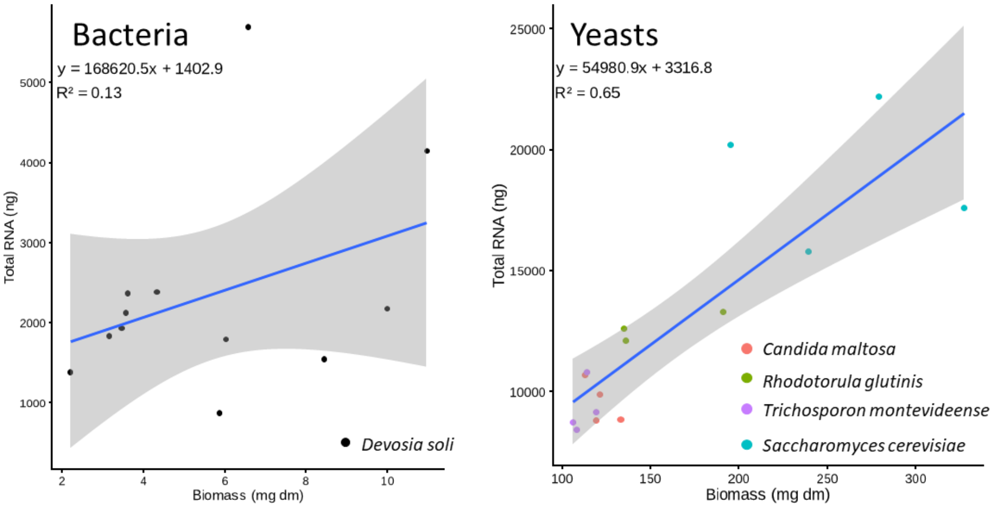

**Fig. S1 | RNA-to-Biomass ratios** for the bacteria *Devosia soli* and four yeast species (see legend). Linear regression suggests a 168621:54981 relationship between bacteria and yeast, supporting the 1:3 scaling difference in ribosome-to-biomass ratio applied in the manuscript between bacteria and fungi.

*Fig. S2 – Relative proportion of bacterial phyla.*

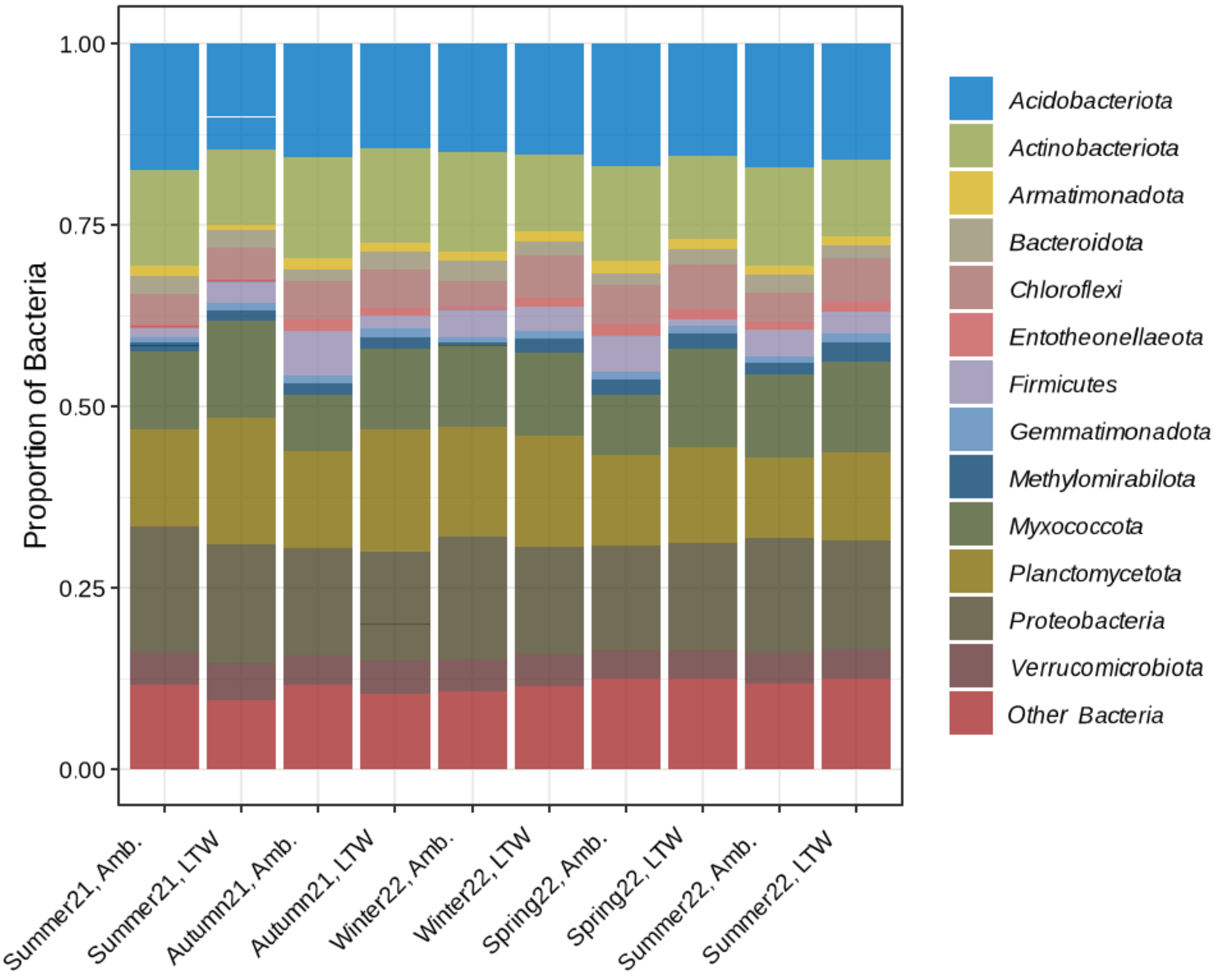

**Fig. S2 | Relative proportion of bacterial phyla in the TOLseq community profiles** (i.e., annotated SSU rRNA transcripts). Summarized for sampling time point (Summer 2021, Autumn 2021, Winter 2022, Spring 2022 and Summer 2022) and soil temperature condition; ambient 'Amb.' and long-termed-warmed 'LTW'. Bacterial members represent a broad range of sizes and life strategies, with both relatively large and slow growing bacteria e.g., *Planctomycetota* (mainly the family *Gemmataceae*) and predatory bacteria (*Myxococcota*, mainly the order *Polyangia*) making up approximately 20% of the *Bacteria* in the community. Contrary, very small bacteria, e.g., *Actinomycota* (*Microtrichales* and *Gaiellales*), *Acidobacteriota* (*Acidobacteriales* and *Solibacteriales*) and *Methylomirabilota* (*Rokubacteriales*) were also abundant (approx. 10%, Data S1).

*Fig. S3 – Dynamics of Ribosome-to-biomass ratio*

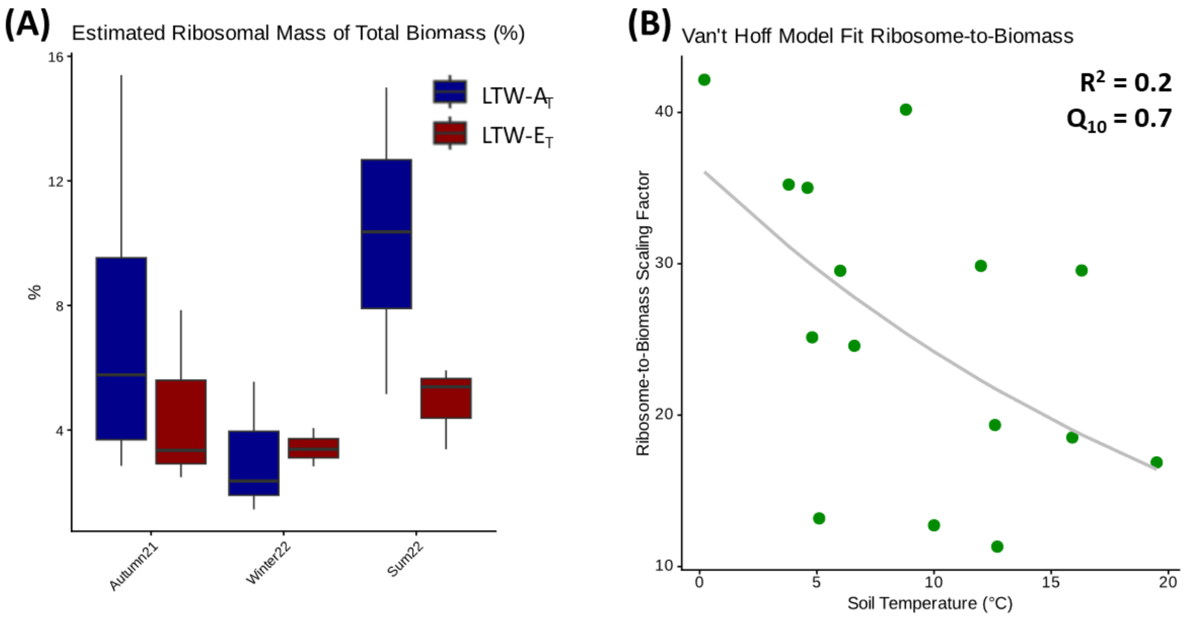

**Fig. S3 | Dynamics of ribosomal mass in relation to seasonal variation and soil temperature.** (**A**) Estimated ribosomal mass as a percentage of total biomass across three sampling periods (Autumn 2021, Winter 2022, and Summer 2022) for ambient '$LTW\text{-}A_T$' (blue) and warmed '$LTW\text{-}E_T$' (red) conditions. Boxplots indicate the median and interquartile range, with whiskers representing the full range of the data. (**B**) Relationship between the ribosome-to-biomass scaling factor and soil temperature (°C). The grey line represents the fit of the Van't Hoff model ($R^2 = 0.2$) with the temperature coefficient, $Q_{10} = 0.7$.

*Fig. S4 – Microbial biomass carbon (MBC) turnover*

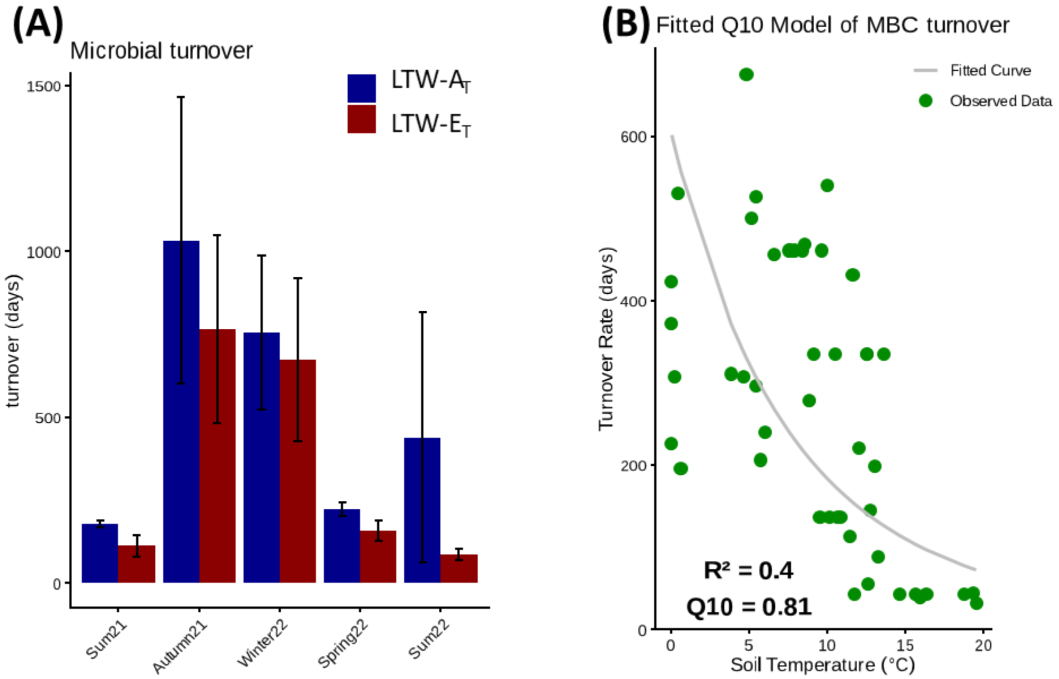

**Fig. S4 | Microbial biomass carbon (MBC) turnover.** (**A**) Mean microbial turnover time (days) across a seasonal gradient for ambient (blue) and warmed (red) plots. Error bars represent ± SD. (**B**) Relationship between microbial turnover rate and soil temperature (°C), fitted with a Van't Hoff model ($R^2 = 0.4$) with the temperature coefficient, $Q_{10} = 0.8$ are shown, indicating a faster turnover as soil temperature increases.

*Fig. S5 – Temperature sensitivity of soil heterotrophic respiration*

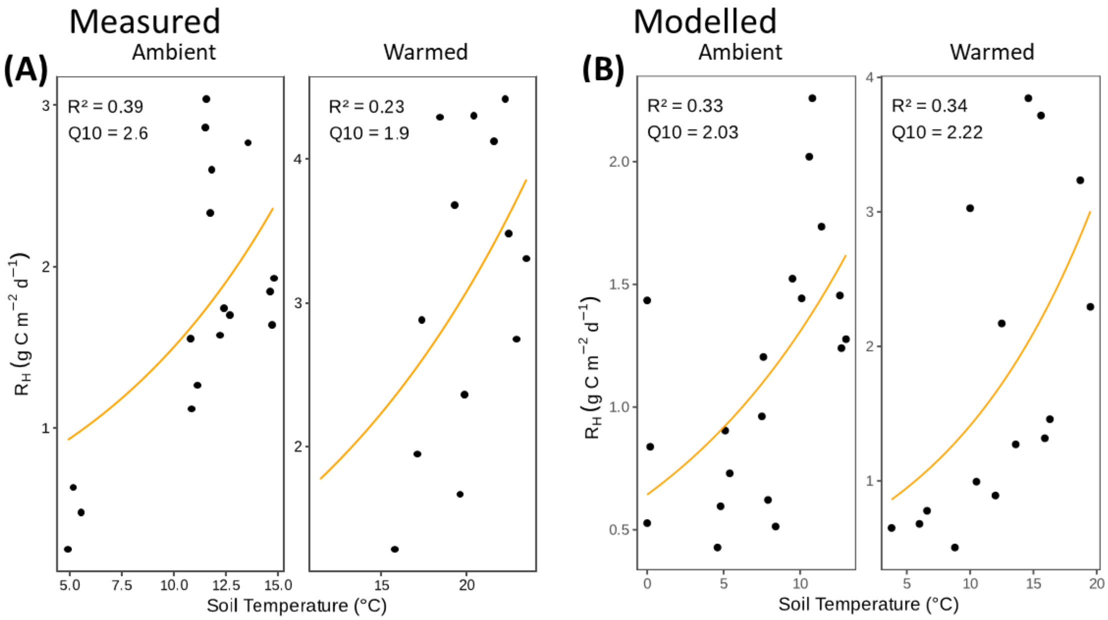

**Fig. S5 | Temperature sensitivity of modelled (A) and measured (B) heterotrophic respiration ($R_H$)** for 'Ambient' and 'Warmed' soil temperature conditions. Measurements *in situ* were obtained between August 2020 and June 2022; however, no measurements were taken in winter (between October and May). In all panels, orange lines represent the fit of the Van't Hoff model and the model fit ($R^2$) and temperature coefficient ($Q_{10}$) are provided

*Fig. S6 – Model parameter sensitivity test*

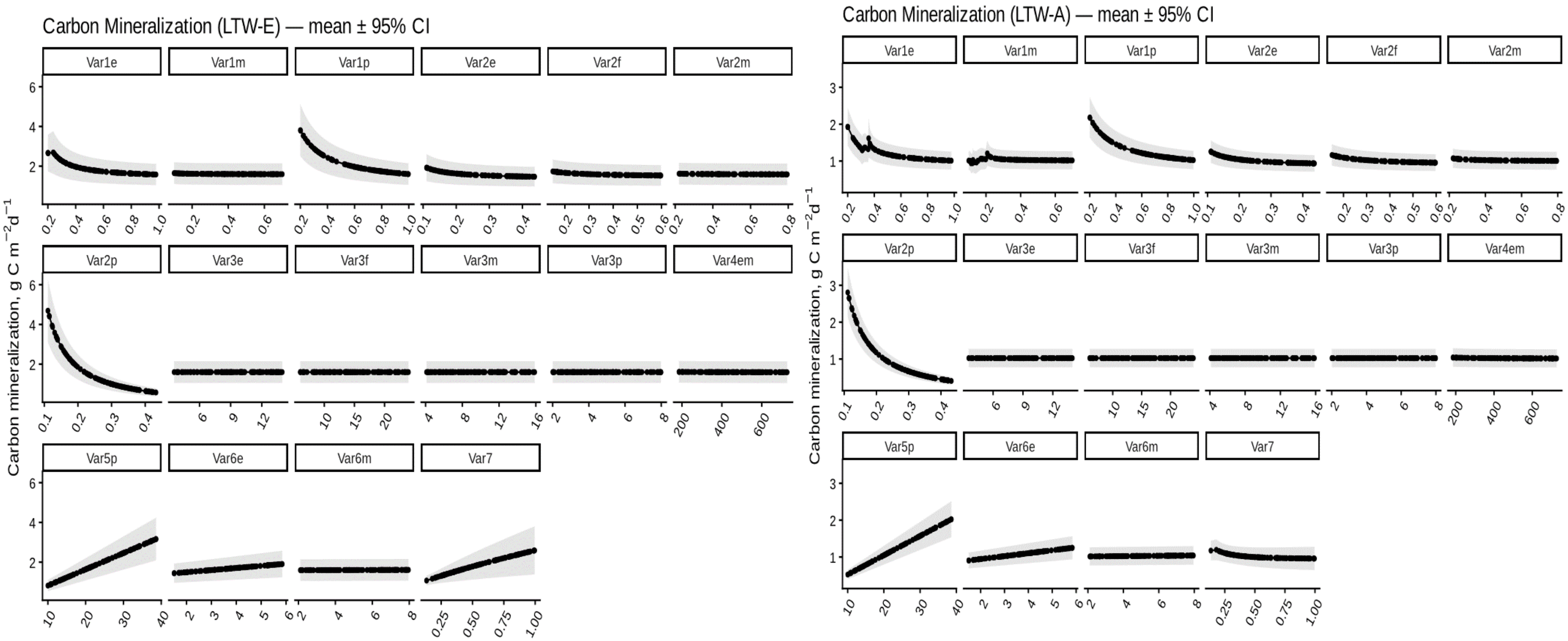

**Fig. S6 | Single-factor sensitivity tests of parameterization for the TOLmodel**. Output calculated for half-to-double variation range in n = 100 permutations per variable and summarized across the data set (n = 3,400; 100 ×34 samples) for the variables (**var**) 'assimilation efficiency' (**1**), 'production efficiency' (**2**), 'C:N ratio' (**3**), 'death rate' (**4**), 'ribosomal-scaling factor for *Prokaryota*' (**5**), 'ribosomes scaling constant' (**6**), 'Q10 for ribosome-to-biomass ratio' (**7**) for prokaryotes (**p**), eukaryotes (**e**) and/or multicellular organisms (**m**) or fungi (**f**). 95% confidence interval is indicated in grey shade. Shown for each soil temperature condition (Long-term warmed (LTW-E), left panel; Ambient (LTW-A), right panel).

*Fig. S7 –Random Forest identification of model parameters*

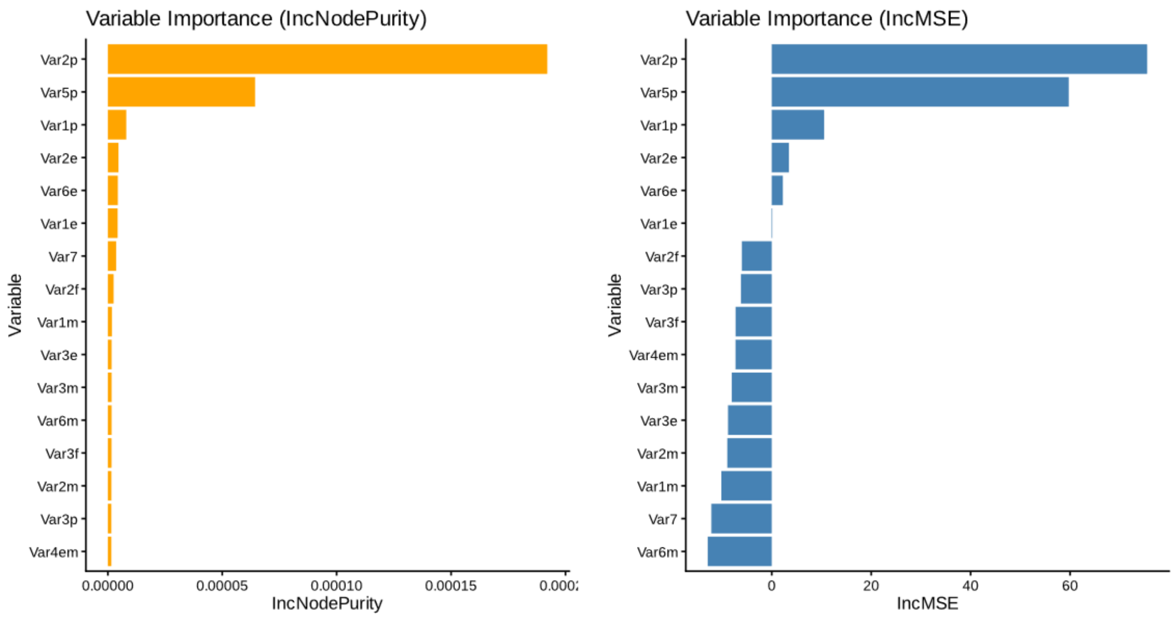

**Fig. S7 | Single-factor sensitivity tests of parameterization for the TOLmodel**. Output calculated for half-to-double variation range in n = 100 permutations per variable and summarized across the data set (n = 3,400; 100 ×34 samples) for the variables (**var**) ‘assimilation efficiency’ (**1**), ‘production efficiency’ (**2**), ‘C:N ratio’ (**3**), ’death rate’ (**4**), ‘ribosomal-scaling factor for *Prokaryota*’ (**5**), ‘ribosomes scaling constant’ (**6**), ‘Q10 for ribosome-to-biomass ratio’ (**7**) for prokaryotes (**p**), eukaryotes (**e**) and/or multicellular organisms (**m**) or fungi (**f**).

*Fig. S8 – Multifactorial sensitivity test.*

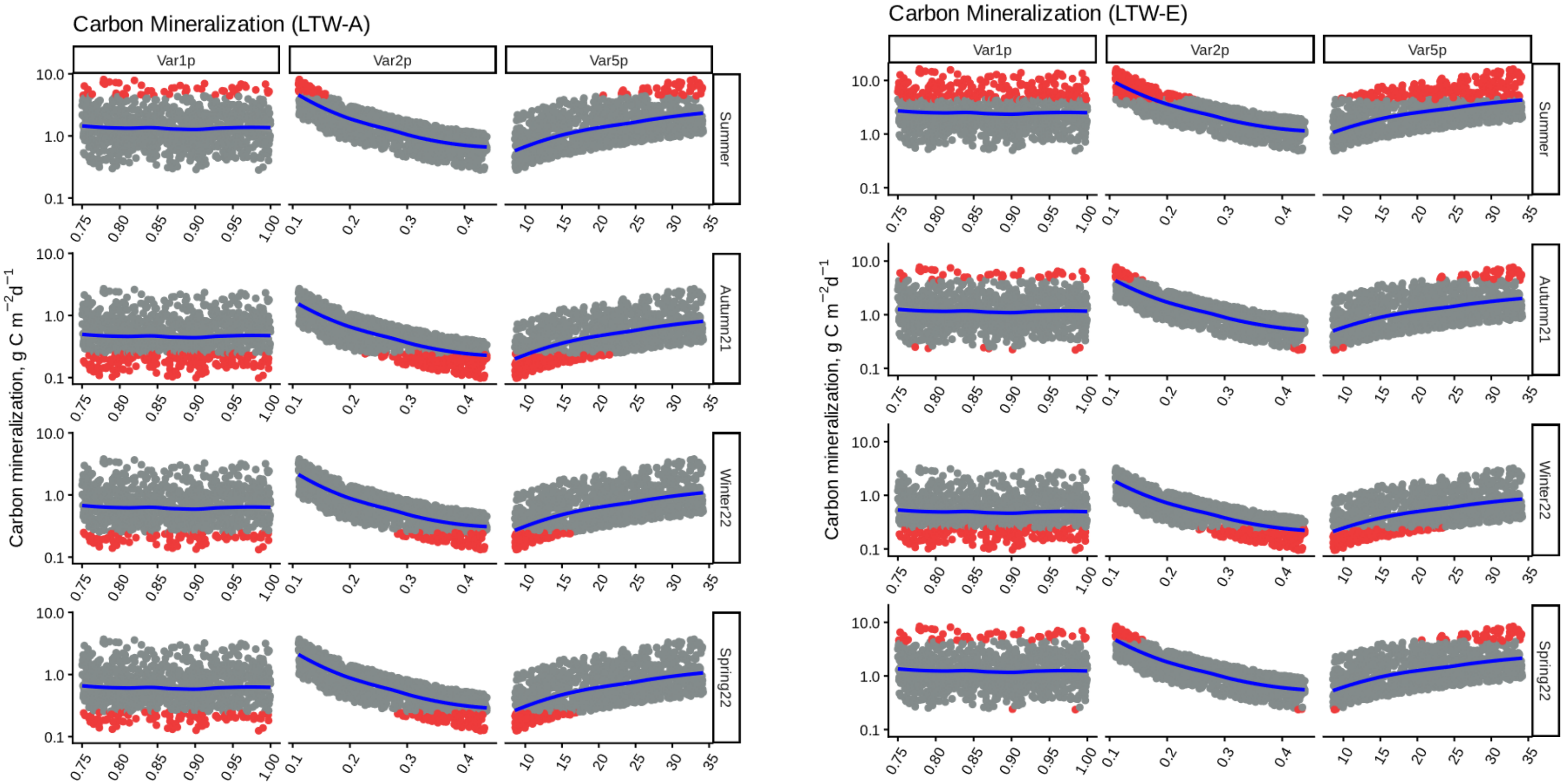

**Fig. S8. | Multifactorial sensitivity tests of parameterization for the TOLmodel**. Output calculated for half-to-double variation range in n = 1,000 permutations per for the variables (**var**) identified as key drivers of the model output (see Fig. S6 and S7): ‘assimilation efficiency’ (**1**), ‘production efficiency’ (**2**), ‘ribosomal-scaling factor for *Prokaryota*’ (**5**), for prokaryotes (**p**). Shown for each soil temperature condition ambient, (LTW-A, left panel) and warmed (LTW-E, right), shown for each season (rows). **Grey**: SOC mineralization rates falling within the range of measured $R_H$; **Red**: values falling outside range of measured $R_H$; **Blue**: LOESS regression.

*Fig. S9 – RNA virus abundance*

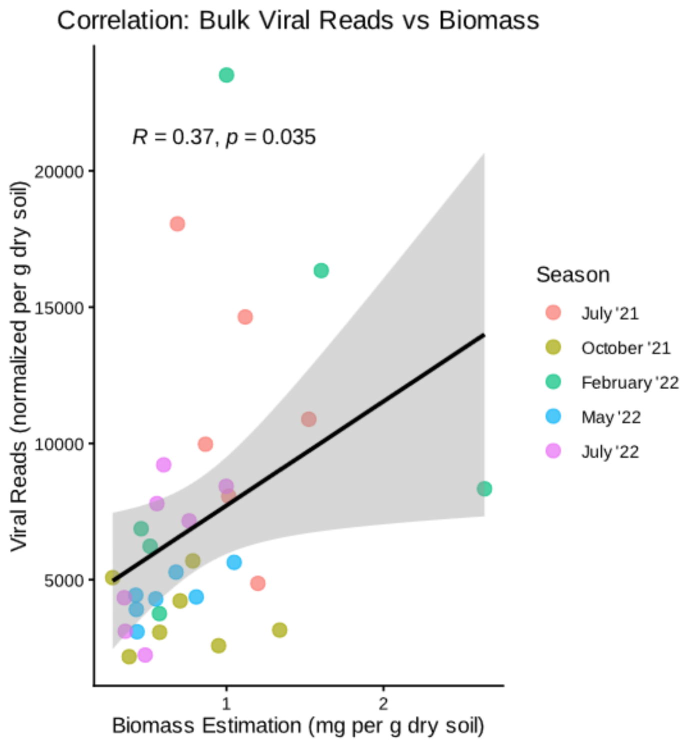

**Fig. S9 | Correlation between soil biomass and viral reads.** Scatter plot showing the relationship between biomass estimation (mg $g^{-1}$ dm soil) and normalized viral reads (reads $g^{-1}$ dm soil), color-coded by sampling season. The solid line represents the linear regression, and the shaded area indicates the 95% confidence interval. A significant positive correlation was observed ($R = 0.37$, $p = 0.035$).

*Fig. S10 – Log2FC in soil heterotrophic respiration between ambient and warmed conditions*

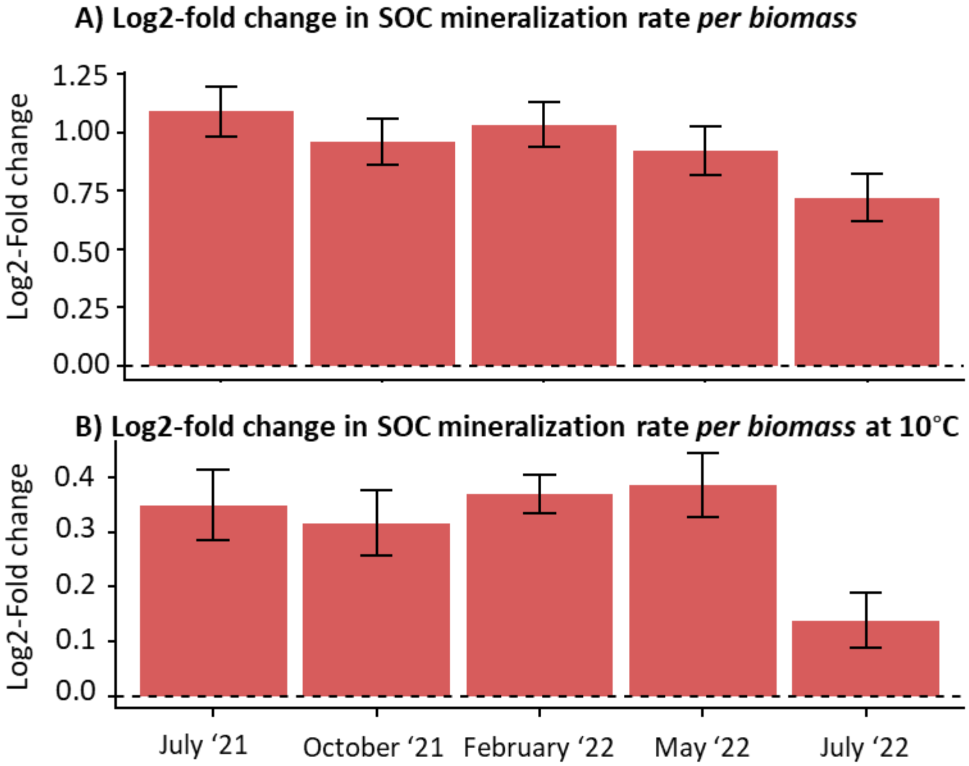

**Fig. S10 | Seasonal variation in SOC mineralization rates.** (**A**) Log2-fold change in mineralization rates (Warmed:Ambient) across five sampling dates from July 2021 to July 2022. (**B**) Log2-fold change measured at a standardized temperature of 10°C. Error bars represent ± SD. Mineralization rates in warmed plots are approximately twofold higher than ambient plots throughout most of the year (log2FC ≈ 1, A); following temperature correction, rates remain significantly elevated by approximately 20–30% (log2FC ≈ 0.35, B).

*Fig. S11 – rrn operons*

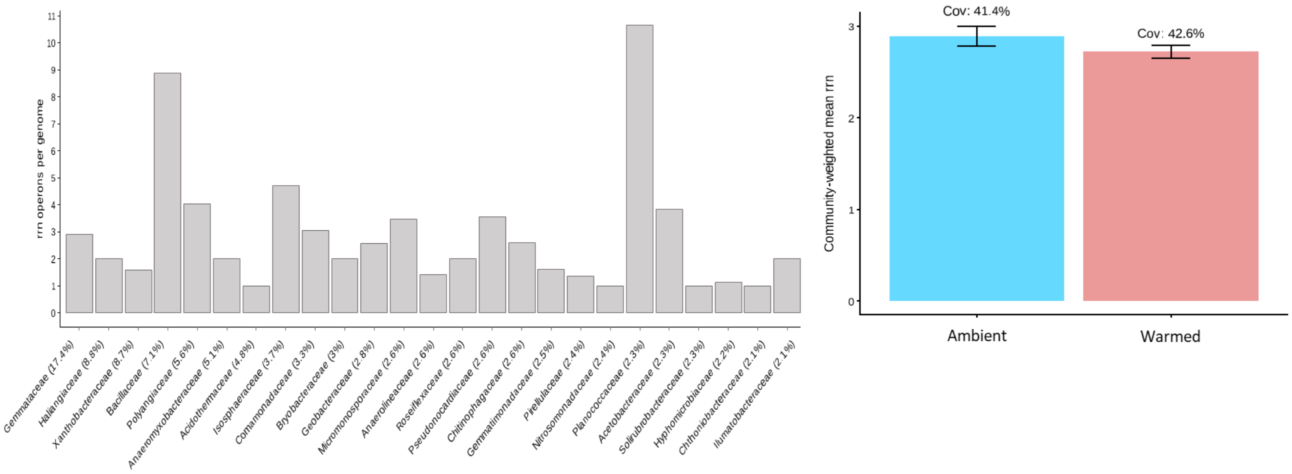

**Fig. S11 | Genomic potential for rapid growth as indicated by *rrn* operon copy number**. (**A**) Distribution of *rrn* operon counts across bacterial families; x-axis labels indicate the family name and its corresponding relative abundance within the community. Higher copy numbers indicate a higher potential for rapid growth. (**B**) Mean *rrn* operon copy number in the Ambient (blue) and Warmed (red) treatments. Error bars represent ± SD and the label indicate the classification coverage.

*Fig. S12 –Richness of Bacteria and Eukaryota in response to warming.*

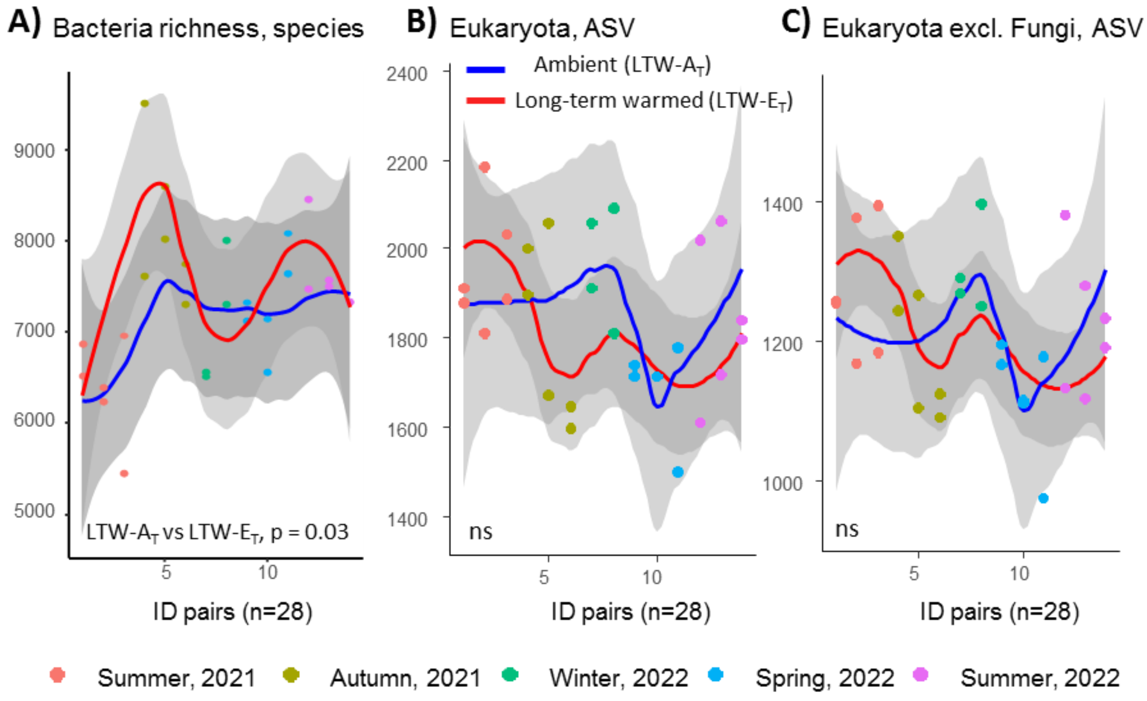

**Fig. S12 | Microbial richness across sampling time points obtained from TOLseq.** Transects are shown in pairs (incomplete pairs excluded). Solid lines indicate LOESS regressions. (**A**) Bacterial richness at the species level. (**B**) Eukaryotic richness at the ASV level, and (**C**) excluding ASVs assigned to Fungi. Reported p-value are from t-tests.

*Fig. S13 – Correlations between the contribution to SOC mineralization of base-level trophic groups and the abundance of higher trophic predators*

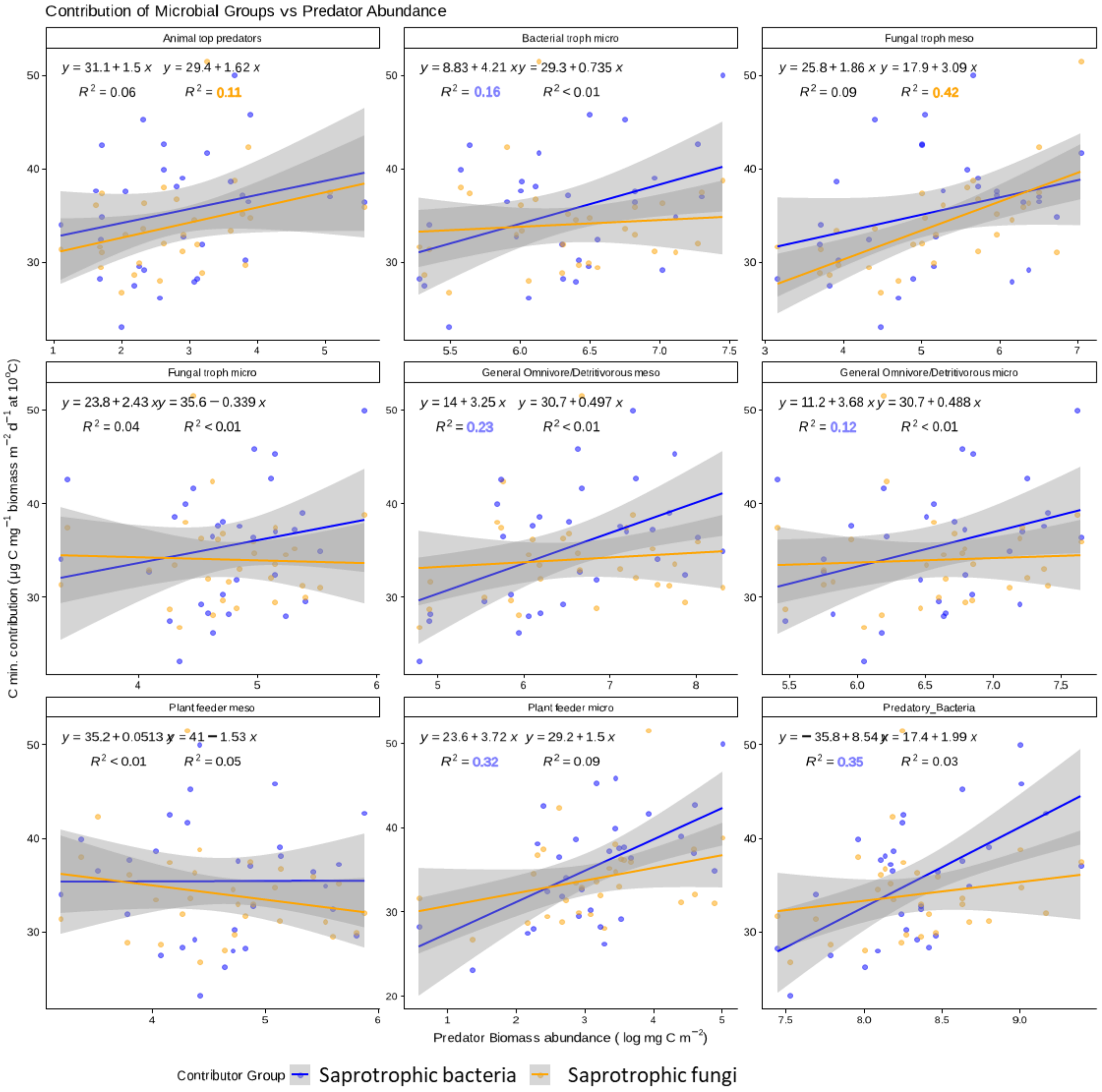

**Fig. S13 | Correlation between predator abundance and the contribution of base-level trophic groups to SOC mineralization.** The panels show the relationship between the log-transformed biomass abundance of nine different predator groups (x-axis) and the temperature normalized carbon mineralization contribution (µg C $mg^{-1}$ biomass $m^{-2}$ $d^{-1}$ at 10°C) of two contributor groups (y-axis): saprotrophic bacteria (blue) and fungi (orange). Each panel contains linear regression lines with their corresponding equations and coefficients of determination ($R^2$), reported $R^2$ in the main text are highlighted. The shaded areas represent the 95% confidence intervals. Ambient winter samples (n=3) were excluded in all plots, since the biomasses at sub-zero temperatures seems to be affected by physiological responses not fully understood.

*Fig. S14 – Soil temperature, showing how transect 3 has cooled down.*

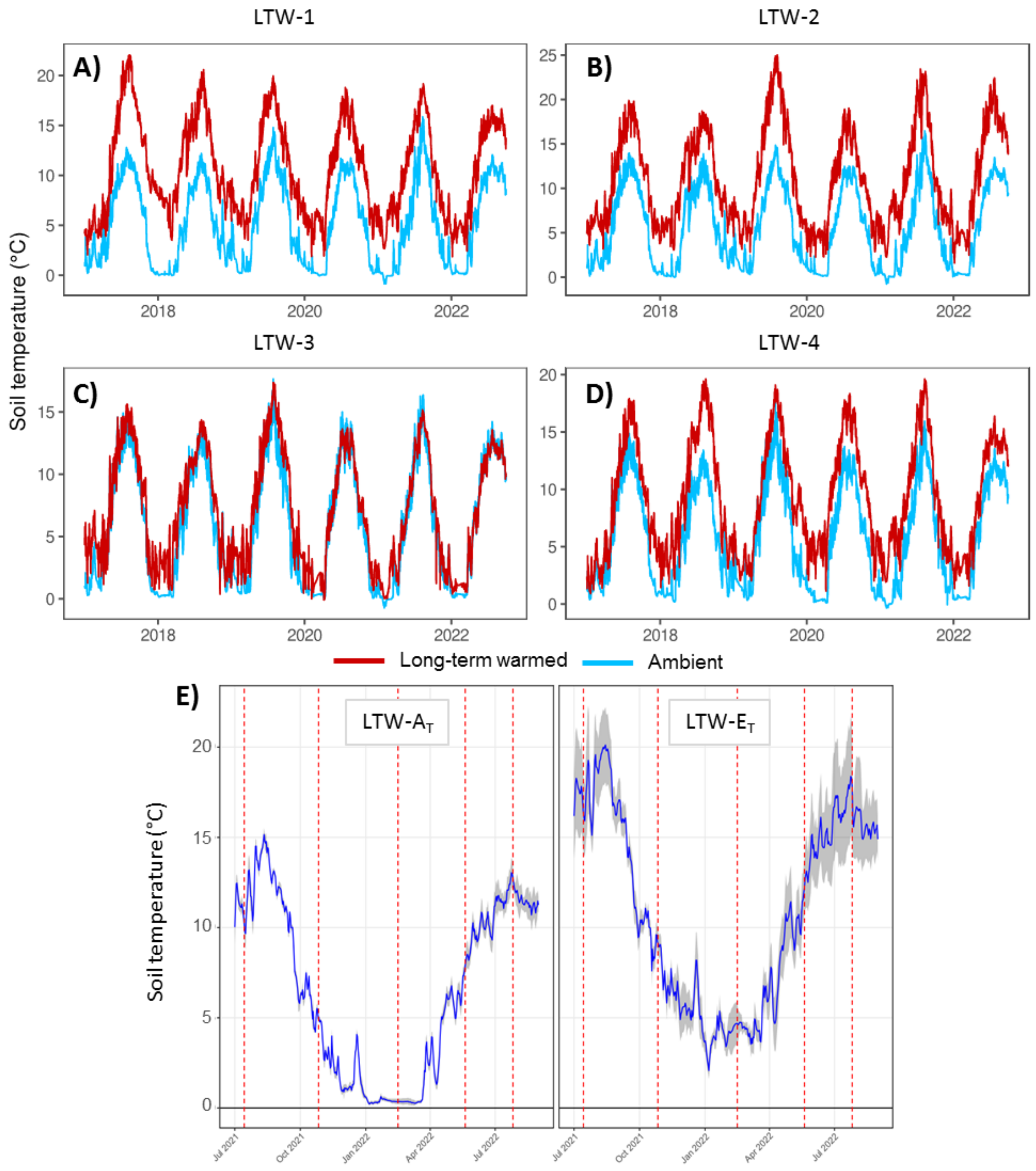

**Fig. S14 | Daily soil temperature under ambient (LTW-A$_T$) and long-term warmed (LTW-E$_T$) conditions,** measured in situ at 12 cm depth at each transect 1–4. Data series from 2017–2023 are shown for each transect (**A–D**). It is seen that the warmed site (red) in transect 3 (LTW-3) has cooled down to the level of ambient (blue). (**E**) Soil temperature profiles for the 2021–2022 sampling year; blue lines indicate LOESS regressions and vertical red lines mark sampling days.

*Fig. S15 – histogram of 18S SSU rRNA contigs*

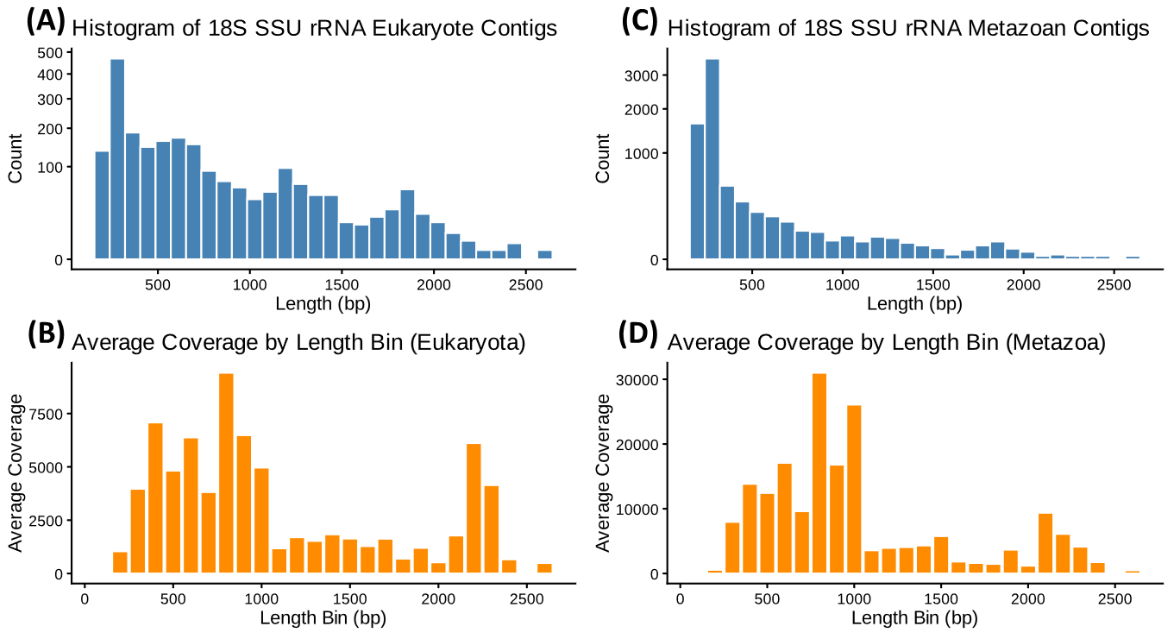

**Fig. S15 | Assembly statistics for 18S SSU rRNA contigs.** (**A, C**) Length distribution of contigs for all eukaryotic reads (**A**) and those assigned to the kingdom *Metazoa* (**C**). Only the longest contig per unique taxonomic annotation is plotted (i.e., showing how many different taxa were able to be assembled to at least X bp); counts are displayed on a square-root scale to highlight distribution across lengths. (**B, D**) Average sequencing coverage across length bins (bp) for all eukaryotes (**B**) and *Metazoa* (**D**), providing a proxy for the relative abundance of these sequences in the dataset.

*Table S1 – TOLmodel parameterization*

**Table S1 | TOLmodel parameters for trophic guilds and carbon pools.**

Food web entities are categorized by organism type (uni_p: unicellular prokaryote; uni_e: unicellular eukaryote; multi: multicellular) and domain (PRO: prokaryote; EUK: eukaryote). Parameters include the assimilation efficiency (a), production effeciency (p), and carbon-to-nitrogen ratio (CN). Functional traits are represented as binary indicators: *isPlant* (plant status), *isDetritus* (detritus status), *canIMM* (capacity for immobilization of nitrogen), and *Detritus Recycling*. Ribosomal mass of a single ribosome (mg) and the associated scaling factor organism type. Abiotic C pools (AtmC: atmospheric carbon; Root: root biomass-C; SOC: soil organic C) are designated as SRA.

| *Trophic guild* | *Org type* | *Domain* | *a*[1] | *p*[2] | *CN*[1] | *isPlant*[3] | *isDetritus*[3] | *canIMM*[3] | *Detritus Recycling*[3] | *One ribosome (mg)* | *Ribosome scaling factor* |
|---|---|---|---|---|---|---|---|---|---|---|---|
| *Archaea* | uni_p | PRO | 1 | 0.22 | 4 | 0 | 0 | 0 | 0 | 4.15E-15 | 24 |
| *Saprotrophic bacteria* | uni_p | PRO | 1 | 0.22 | 4 | 0 | 0 | 1 | 0 | 4.15E-15 | 24 |
| *Predatory_Archaea* | uni_p | PRO | 1 | 0.22 | 4 | 0 | 0 | 0 | 0 | 4.15E-15 | 24 |
| *Predatory_Bacteria* | uni_p | PRO | 1 | 0.22 | 4 | 0 | 0 | 0 | 0 | 4.15E-15 | 24 |
| *Algae* | uni_e | EUK | 1 | 0.22 | 7 | 0 | 0 | 0 | 0 | 5.48E-15 | 72 |
| *Fungi* | uni_e | EUK | 0.9 | 0.3 | 12 | 0 | 0 | 1 | 0 | 5.48E-15 | 72 |
| *Arbuscular mycorrhizal fungi* | uni_e | EUK | 0.9 | 0.3 | 12 | 0 | 0 | 1 | 0 | 5.48E-15 | 72 |
| *Metazoa_TopPredator* | uni_e | EUK | 0.9 | 0.22 | 7 | 0 | 0 | 0 | 0 | 5.48E-15 | 72 |
| *Predator_Fungi* | uni_e | EUK | 0.9 | 0.3 | 12 | 0 | 0 | 0 | 0 | 5.48E-15 | 72 |
| *Prostists_Autotroph* | uni_e | EUK | 1 | 0.22 | 7 | 0 | 0 | 0 | 0 | 5.48E-15 | 72 |
| *Prostists_Bacterivore* | uni_e | EUK | 0.9 | 0.22 | 7 | 0 | 0 | 0 | 0 | 5.48E-15 | 72 |
| *Prostists_Eukaryvore* | uni_e | EUK | 0.9 | 0.22 | 7 | 0 | 0 | 0 | 0 | 5.48E-15 | 72 |
| *Prostists_Mixotrophic* | uni_e | EUK | 0.9 | 0.22 | 7 | 0 | 0 | 0 | 0 | 5.48E-15 | 72 |
| *Prostists_Omnivore* | uni_e | EUK | 0.9 | 0.22 | 7 | 0 | 0 | 0 | 0 | 5.48E-15 | 72 |
| *Prostists_Parasite (Animals)* | uni_e | EUK | 0.9 | 0.22 | 7 | 0 | 0 | 0 | 0 | 5.48E-15 | 72 |
| *Prostists_Parasite (Plants)* | uni_e | EUK | 0.9 | 0.22 | 7 | 0 | 0 | 0 | 0 | 5.48E-15 | 72 |
| *Prostists_Saprotrophic/Osmotrophic* | uni_e | EUK | 0.9 | 0.22 | 7 | 0 | 0 | 0 | 0 | 5.48E-15 | 72 |
| *Rhizaria_animal_parasite* | uni_e | EUK | 0.9 | 0.22 | 7 | 0 | 0 | 0 | 0 | 5.48E-15 | 72 |
| *Rhizaria_autotroph* | uni_e | EUK | 1 | 0.22 | 7 | 0 | 0 | 0 | 0 | 5.48E-15 | 72 |
| *Rhizaria_bacterivore* | uni_e | EUK | 0.9 | 0.22 | 7 | 0 | 0 | 0 | 0 | 5.48E-15 | 72 |
| *Rhizaria_eukaryvore* | uni_e | EUK | 0.9 | 0.22 | 7 | 0 | 0 | 0 | 0 | 5.48E-15 | 72 |

| | | | | | | | | | | | |
|---|---|---|---|---|---|---|---|---|---|---|---|
| *Rhizaria_omnivore* | uni_e | EUK | 0.9 | 0.22 | 7 | 0 | 0 | 0 | 0 | 5.48E-15 | 72 |
| *Rhizaria_plant_parasite* | uni_e | EUK | 0.9 | 0.22 | 7 | 0 | 0 | 0 | 0 | 5.48E-15 | 72 |
| *Metazoa_Collembola_Detriti_FungalFeeders* | multi | EUK | 0.5 | 0.4 | 8 | 0 | 0 | 0 | 0 | 6.14E-15 | 96 |
| *Metazoa_Detritivore_Microbivore* | multi | EUK | 0.5 | 0.4 | 8 | 0 | 0 | 0 | 0 | 6.14E-15 | 96 |
| *Metazoa_Detritivores* | multi | EUK | 0.5 | 0.4 | 8 | 0 | 0 | 0 | 0 | 6.14E-15 | 96 |
| *Metazoa_FungalFeeders* | multi | EUK | 0.5 | 0.4 | 8 | 0 | 0 | 0 | 0 | 6.14E-15 | 96 |
| *Metazoa_Herbivores* | multi | EUK | 0.5 | 0.4 | 8 | 0 | 0 | 0 | 0 | 6.14E-15 | 96 |
| *Metazoa_Parasites (Animal)* | multi | EUK | 0.5 | 0.4 | 8 | 0 | 0 | 0 | 0 | 6.14E-15 | 96 |
| *Nematoda_Bacterial feeding* | multi | EUK | 0.5 | 0.4 | 8 | 0 | 0 | 0 | 0 | 6.14E-15 | 96 |
| *Nematoda_Fungal feeding* | multi | EUK | 0.5 | 0.4 | 8 | 0 | 0 | 0 | 0 | 6.14E-15 | 96 |
| *Nematoda_Omnivore* | multi | EUK | 0.5 | 0.4 | 8 | 0 | 0 | 0 | 0 | 6.14E-15 | 96 |
| *Nematoda_Plant feeding* | multi | EUK | 0.5 | 0.4 | 8 | 0 | 0 | 0 | 0 | 6.14E-15 | 96 |
| *Nematoda_Predators* | multi | EUK | 0.5 | 0.4 | 8 | 0 | 0 | 0 | 0 | 6.14E-15 | 96 |
| *AtmC* | SRA | | 0 | 0 | 0 | 0 | 1 | 0 | 0 | | |
| *Root* | SRA | | 0 | 0 | 0 | 1 | 0 | 0 | 0 | | |
| *SOC* | SRA | | 0 | 0 | 0 | 0 | 1 | 0 | 1 | | |

[1] Adapted from recent application and original publication on soil food web modelling (Buchkowski et al., 2023, Holtkamp 2011, Hunt et al., 1987, de Ruiter 1993). [2] We adapted a carbon-use-efficiency (CUE) for microorganisms of 0.22 determined from the site (walker et al 2018). [3] Following (Buchkowski et al., 2023).

*Table S2 – Trophic guild abundance and contributions to SOC mineralization*

**Table S2 |Seasonal abundance and relative contribution to SOC mineralization of trophic guilds under ambient and warmed conditions.** Values are presented as mean ± standard deviation. Absolute abundance is expressed in grams per square meter ($g^{-1}$ $m^2$), while contributions are expressed as a percentage (%) of the SOC mineralization.

| *Trophic guild* | *Ambient* | | | | *Warmed* | | | |
|---|---|---|---|---|---|---|---|---|
| **Abundance (g m$^2$)** | **Autumn** | **Winter** | **Spring** | **Summer** | **Autumn** | **Winter** | **Spring** | **Summer** |
| *Saprotrophic bacteria* | 34.3 ± 6.4 | 70.3 ± 34.3 | 32.9 ± 11.8 | 37.2 ± 10.3 | 40.9 ± 33.6 | 31.6 ± 4.5 | 29.4 ± 7.5 | 35.2 ± 15.5 |
| *Predatory bacteria* | 3.1 ± 1 | 9.4 ± 4.7 | 3.2 ± 1.4 | 4.7 ± 1.2 | 5.7 ± 5.5 | 4.2 ± 0.4 | 5 ± 2.7 | 5.6 ± 2.7 |
| *Bacterial troph micro* | 0.6 ± 0.4 | 1.9 ± 1.1 | 0.4 ± 0.3 | 0.8 ± 0.3 | 0.6 ± 0.8 | 0.5 ± 0.1 | 0.5 ± 0.2 | 0.8 ± 0.6 |
| *Fungi* | 4.8 ± 1 | 12.9 ± 4.9 | 4 ± 2.3 | 8.3 ± 5.5 | 3.3 ± 4.8 | 2.7 ± 0.7 | 3.5 ± 0.4 | 5.6 ± 3.4 |
| *Fungal troph micro* | 0.1 ± 0 | 0.4 ± 0.3 | 0.1 ± 0.1 | 0.2 ± 0.1 | 0.1 ± 0.1 | 0.1 ± 0 | 0.1 ± 0 | 0.1 ± 0.1 |
| *Fungal troph meso* | 0.3 ± 0.3 | 2 ± 2.1 | 0.1 ± 0.1 | 0.5 ± 0.3 | 0.2 ± 0.4 | 0.1 ± 0.1 | 0.3 ± 0.3 | 0.2 ± 0.1 |
| *Arbuscular mycorrhizal fungi* | 0.6 ± 0.1 | 1.9 ± 0.7 | 0.7 ± 0.5 | 1.7 ± 1.3 | 0.5 ± 0.7 | 0.5 ± 0.2 | 0.7 ± 0.4 | 1.5 ± 1.4 |
| *Autotrophs* | 0.1 ± 0 | 0.3 ± 0.1 | 0.1 ± 0.1 | 0.1 ± 0.1 | 0.1 ± 0.1 | 0 ± 0 | 0.1 ± 0 | 0.1 ± 0.1 |
| *Plant feeder micro* | 0.05 ± 0.04 | 0.02 ± 0.05 | 0.08 ± 0.11 | 0.01 ± 0.06 | 0.06 ± 0.01 | 0.04 ± 0.01 | 0.02 ± 0.01 | 0.03 ± 0.01 |
| *Plant feeder meso* | 0.2 ± 0.08 | 0.09 ± 0.07 | 0.22 ± 0.05 | 0.16 ± 0.12 | 0.11 ± 0.02 | 0.05 ± 0.12 | 0.08 ± 0.05 | 0.08 ± 0.07 |
| *General Omnivore/Detritivore micro* | 0.8 ± 0.4 | 2.7 ± 1.3 | 0.6 ± 0.3 | 1.2 ± 0.5 | 0.6 ± 0.7 | 0.6 ± 0.2 | 0.6 ± 0.3 | 1 ± 0.6 |
| *General Omnivore/Detritivore meso* | 0.4 ± 0.2 | 2.6 ± 3.2 | 0.2 ± 0.1 | 2 ± 1.2 | 1.1 ± 1 | 0.7 ± 0.2 | 0.5 ± 0.2 | 1.1 ± 0.8 |
| *Animal top predators* | 0.05 ± 0.09 | 0.01 ± 0.09 | 0.01 ± 0.01 | 0.02 ± 0.01 | 0.02 ± 0.01 | 0.05 ± 0.02 | 0.02 ± 0 | 0.03 ± 0.03 |
| *Animal predators micro* | 0.12 ± 0.05 | 0.07 ± 0.1 | 0.27 ± 0.23 | 0.07 ± 0.11 | 0.12 ± 0.04 | 0.07 ± 0.05 | 0.04 ± 0.01 | 0.05 ± 0.01 |
| **Contributions (%)** | | | | | | | | |
| *Saprotrophic bacteria* | 84 ± 2 | 83 ± 2 | 86 ± 2 | 81 ± 3 | 88 ± 5 | 86 ± 1 | 84 ± 1 | 84 ± 2 |
| *Predatory bacteria* | 5.3 ± 0.5 | 6.9 ± 0.4 | 5.7 ± 0.3 | 6.6 ± 0.7 | 7.2 ± 0.5 | 7.1 ± 0.6 | 7.8 ± 1.2 | 7.6 ± 0.6 |
| *Bacterial troph micro* | 1 ± 0.3 | 1.3 ± 0.2 | 0.7 ± 0.2 | 1 ± 0.1 | 0.5 ± 0.5 | 0.9 ± 0.1 | 0.8 ± 0.1 | 1 ± 0.4 |
| *Fungi* | 6.9 ± 1.3 | 6.6 ± 1.9 | 5.2 ± 1.1 | 7.7 ± 2.4 | 2.4 ± 2.4 | 3.6 ± 0.7 | 4.9 ± 1.5 | 5 ± 1.6 |
| *Fungal troph micro* | 0.19 ± 0.1 | 0.27 ± 0.1 | 0.23 ± 0.1 | 0.21 ± 0.05 | 0.06 ± 0.1 | 0.15 ± 0.05 | 0.19 ± 0.03 | 0.17 ± 0.1 |

| | | | | | | | | |
|---|---|---|---|---|---|---|---|---|
| *Fungal troph meso* | 0.2 ± 0.12 | *NA* | 0.04 ± 0.02 | 0.14 ± 0.12 | 0.07 ± 0.09 | 0.08 ± 0.05 | 0.13 ± 0.12 | 0.05 ± 0.03 |
| *Arbuscular mycorrhizal fungi* | 0.9 ± 0.2 | 1 ± 0.4 | 0.8 ± 0.3 | 1.6 ± 0.6 | 0.4 ± 0.4 | 0.7 ± 0.3 | 0.9 ± 0.2 | 1.2 ± 0.6 |
| *Autotrophs* | 0.12 ± 0.04 | 0.19 ± 0.05 | 0.14 ± 0.06 | 0.17 ± 0.1 | 0.08 ± 0.07 | 0.08 ± 0.04 | 0.1 ± 0.03 | 0.1 ± 0.05 |
| *Plant feeder micro* | 0.03 ± 0.02 | 0.04 ± 0.04 | 0.02 ± 0.01 | 0.05 ± 0.03 | 0.03 ± 0.03 | 0.03 ± 0.01 | 0.04 ± 0.02 | 0.05 ± 0.04 |
| *Plant feeder meso* | 0.1 ± 0.03 | *NA* | 0.13 ± 0.06 | 0.11 ± 0.05 | 0.02 ± 0.02 | 0.07 ± 0.04 | 0.05 ± 0.01 | 0.04 ± 0.03 |
| *General Omnivore/Detritivore micro* | 1 ± 0.3 | 0.9 ± 0 | 0.8 ± 0.3 | 1.1 ± 0.3 | 0.5 ± 0.4 | 0.8 ± 0.3 | 0.8 ± 0.2 | 0.9 ± 0.3 |
| *General Omnivore/Detritivore meso* | 0.3 ± 0.1 | *NA* | 0.2 ± 0.1 | 0.5 ± 0.2 | 0.6 ± 0.8 | 0.5 ± 0.1 | 0.2 ± 0.1 | 0.3 ± 0.1 |
| *Animal top predators* | 0.02 ± 0.01 | *NA* | 0.03 ± 0.03 | 0.04 ± 0.04 | 0.04 ± 0.06 | 0.03 ± 0.01 | 0.04 ± 0.02 | 0.03 ± 0.03 |
| *Animal predators micro* | 0.1 ± 0.04 | 0.13 ± 0.06 | 0.09 ± 0.05 | 0.12 ± 0.03 | 0.05 ± 0.05 | 0.06 ± 0.01 | 0.07 ± 0.01 | 0.11 ± 0.06 |

*Table S3 – Grassland I, II and III contributors*

**Table S3 | Relative contribution to SOC mineralization of trophic guilds in Grassland I, II and III.** Values are presented as mean ± SD specified for each season; summer (Sum), autumn (Aut), Winter, spring.

| *Trophic Guild* | *Grassland I* | | | | *Grassland II* | | | | *Grassland III* | | | |
|---|---|---|---|---|---|---|---|---|---|---|---|---|
| | **Sum** | **Aut** | **Winter** | **Spring** | **Sum** | **Aut** | **Winter** | **Spring** | **Sum** | **Aut** | **Winter** | **Spring** |
| *Saprotrophic bacteria* | 81 ± 17 | 78 ± 16 | 72 ± 12 | 81 ± 34 | 75 ± 25 | 73 ± 17 | 73 ± 1 | 68 ± 39 | 81 ± 77 | 81 ± 83 | 76 ± 45 | 83 ± 81 |
| *Predatory bacteria* | 5 ± 1 | 5 ± 1 | 5 ± 1 | 5 ± 2 | 4 ± 2 | 6 ± 2 | 5 ± 0 | 4 ± 2 | 4 ± 5 | 3 ± 2 | 3 ± 5 | 4 ± 3 |
| *Archaea* | 3 ± 1 | 3 ± 2 | 2 ± 1 | 3 ± 1 | 4 ± 2 | 4 ± 1 | 4 ± 1 | 6 ± 11 | 0.6 ± 0.9 | 0.5 ± 0.3 | 0.6 ± 0.3 | 1 ± 1 |
| *Fungi* | 6 ± 1 | 5 ± 2 | 10 ± 4 | 6 ± 3 | 7 ± 3 | 6 ± 1 | 7 ± 1 | 11 ± 14 | 2 ± 2 | 2 ± 3 | 7 ± 3 | 0.9 ± 0.8 |
| *Arbuscular mycorrhizal* | 0.9 ± 0.2 | 0.9 ± 0.5 | 2 ± 0.4 | 0.9 ± 0.8 | 0.7 ± 0.4 | 1 ± 1 | 2 ± 0.6 | 2 ± 1.3 | 3 ± 4 | 4 ± 1 | 3 ± 2 | 1 ± 2 |
| *Prostists omnivore* | 2 ± 0.3 | 2 ± 0.6 | 2 ± 0.5 | 2 ± 0.6 | 4 ± 1 | 4 ± 2 | 2 ± 0.4 | 4 ± 2 | 6 ± 5 | 7 ± 8 | 4 ± 4 | 6 ± 8 |
| *Prostists bacterivore* | 2 ± 0.2 | 2 ± 0.5 | 2 ± 0.4 | 2 ± 1 | 2 ± 1 | 3 ± 1 | 2 ± 0.3 | 2 ± 2 | 3 ± 5 | 1 ± 2 | 2 ± 3 | 2 ± 2 |
| *Rhizaria bacterivore* | — | — | 0.5 ± 0.1 | — | — | 0.5 ± 0.2 | 0.6 ± 0.1 | 0.5 ± 0.5 | 0.1 ± 0.1 | 0.04 ± 0 | 0.2 ± 0.7 | 0.02 ± 0 |
| *Rhizaria omnivore* | — | — | — | — | — | — | — | 0.5 ± 0.5 | 0.1 ± 0.3 | 0.1 ± 0.1 | 0.1 ± 0.2 | 0.2 ± 0.2 |
| *Metazoa collembola* | 0.4 ± 0.2 | 1 ± 0.5 | 2 ± 0.3 | 0.4 ± 0.2 | 0.7 ± 0.3 | 1.3 ± 0.7 | 3 ± 0.2 | 0.6 ± 0.7 | — | — | — | — |
| *Metazoa fungal feeders* | — | — | — | — | — | — | 0.5 ± 0.1 | — | 0.1 ± 0.1 | 0.1 ± 0.1 | 0.3 ± 0.5 | 0.2 ± 0.6 |
| *Nematoda plant feeding* | — | — | — | — | 0.4 ± 0.1 | — | — | — | 0.1 ± 0.2 | 0.01 ± 0.02 | — | 0.1 ± 0.2 |

*Table S4 – Daily weather conditions July 2021.*

**Table S4 | Daily weather conditions during *in situ* soil $CO_2$ efflux measurements in July 2021**. TOLseq soil samples were collected on 14 July (in bold). Values are daily means ± SD unless otherwise noted. *Tair*: daily air temperature; *VPD*: daily vapor pressure deficit; *$VPD_{max}$*: daily maximum vapor pressure deficit; *Solar radiation*: daily integrated shortwave radiation; *7-day rainfall total*: cumulative precipitation during the previous 7 days.

| **Date** | **$T_{air}$ (°C)** | **VPD (kPa)** | **$VPD_{max}$ (Kpa)** | **Solar radiation (MJ $m^{-2}$ $d^{-1}$)** | **7-day rainfall total (mm)** |
|---|---|---|---|---|---|
| 07/07/2021 | 10.11 ± 1.20 | 0.12 ± 0.11 | 0.41 | 12.77 | 1.3 |
| **14/07/2021** | 9.14 ± 0.51 | 0.10 ± 0.07 | 0.27 | 6.14 | 19.1 |
| 28/07/2021 | 14.56 ± 3.69 | 0.51 ± 0.28 | 1.15 | 21.07 | 107.4 |

*Data S1 – Abundance table of SSU rRNA annotated taxa (.csv)*

Normalized read counts (to 1 million) of annotated SSU rRNA. A total of 30.191 unique taxonomic annotations (rows). Column A ("Taxonomy") provide the full LCA taxon string followed by each sample in columns J to AW (n = 40). Sample IDs are listed in Data S4.

*Data S2 – Soil food web interaction (feeding) matrix (.csv)*

Interaction matrix (row eats column) of the 34 identified soil food web entities.

*Data S3 – Trophic classification (.csv)*

Abundance table of each member of the trophic nodes.

*Data S4 – Sample metadata (.xlsx)*

List of sample IDs and corresponding metadata for each sample.
